# Discovery and characterization of FX-909, a covalent inverse agonist of PPARG rationally designed to impose a powerful repressive bias in PPARG for the treatment of PPARG/RXRA-activated muscle-invasive urothelial cancers

**DOI:** 10.1101/2025.05.08.652848

**Authors:** Jacob I. Stuckey, Jennifer A. Mertz, Jonathan E. Wilson, Kaylyn E. Williamson, Yong Li, Miljan Kuljanin, Byron DeLaBarre, Gregg Chenail, Phuong A. Nguyen, Christopher M. Bailey, William W. Motley, James E. Audia, Robert J. Sims

## Abstract

We report our mechanistic investigation into the conformationally-driven ‘activation bias’ of PPARG in muscle-invasive urothelial cancer (MIUC) and our efforts to pharmacologically reverse this activation bias through covalent PPARG inverse agonism. We utilized studies into tumor-associated mutations in both PPARG and RXRA, as well as a combination of structure-based drug design merged with insights from biochemical mechanistic studies to discover FX-909, a first-in-class clinical PPARG inverse agonist that robustly enforces a conformationally ‘repressive’ state of PPARG, even in highly activated contexts such as RXRA S427F mutation and PPARG amplification. FX-909 is a potent, highly selective, and powerful suppressor of PPARG transcriptional activity through enhancement of PPARG nuclear corepressor binding (NCOR) affinity. Treatment with FX-909 resulted in selective growth inhibition in PPARG-activated MIUC cell lines. Further, FX-909 achieved durable regressions in xenograft models of MIUC through inverse agonism of PPARG. FX-909 is the first chemical tool available to the community that is capable of recapitulating PPARG genetic knockout *in vivo* and is currently in clinical development for the treatment of intractable MIUC.

## Intro/Background

Peroxisome proliferator-activated receptor gamma (PPARG) is a nuclear hormone receptor and transcription factor that is most well understood as a physiological fatty acid sensor regulating lipid homeostasis and energy balance^1^. Additionally, PPARG plays an essential, lineage determining role in the luminal cell layers of the normal urothelium^2,3^ and is crucial for its homeostasis and regeneration^3^. Transcriptional activity of PPARG is regulated through the conformation of helix 12 (H12), located at the C-terminus of the protein^4^. For isolated PPARG, H12 exhibits conformational mobility, favoring the binding of CRNR box motifs found in co-repressors^4,5^. H12 is biased towards an active conformation in response to ligands, resulting in a high affinity conformational state for LxxLL motifs present in various co-activators^4^. In cells, DNA binding and activation of PPARG requires heterodimerization with retinoid X receptor alpha (RXRA)^6^.

PPARG drives the initiation and development of luminal lineage urothelial cancer (UC) and is the top selective dependency factor for UC cell lines in genome-wide screens^7–10^. Genetic evidence of recurrent activation of PPARG transcription and signaling suggests a functional dependency in UC^11^. Genetic profiling of UC patients has identified recurrent alterations in the PPARG gene, including focal amplification, missense mutations, and gene fusions. Missense mutations in the heterodimeric binding partner of PPARG, RXRA, have also been observed^11–13^. RXRA^S427F^ has previously been shown to drive activation of PPARG through enhanced interaction with PPARG and by increasing stabilization of the active conformation of H12^12,14^. The ability of missense mutations in PPARG to drive an activated state is less clear but molecular dynamics simulations paired with <u>h</u>ydrogen <u>d</u>euterium e<u>x</u>change (HDX) studies have indicated these mutations also stabilize the active conformation of H12 in PPARG^12^.

T0070907 was the first PPARG inverse agonist described in 2002^15^. T0070907 is a covalent modifier of PPARG at cysteine 313 (C313) that induces a repressive conformational landscape that enhances the affinity of PPARG for co-repressor peptides while decreasing the affinity for co-activator peptides. The conformational mechanism underlying T0070907’s inverse agonism was elucidated by X-ray crystallography in 2020^16^. The mechanism involves induction of a conformational state in which H12 inserts into the ligand binding pocket of PPARG to form distinct interactions with the residual covalent adduct of T0070907. Noncovalent inverse agonists have also been described for PPARG which do not display this conformational state of H12 and have historically exhibited weaker recruitment of co- repressor peptides relative to T0070907^11,14^. Recent work has produced additional PPARG inverse agonists, both covalent and noncovalent, each with reported abilities to recruit co-repressor peptides similar to T0070907^14,17^. Covalent inverse agonists with modestly enhanced co-repressor recruitment were also recently described with a preference for recruitment of NCOR2 over NCOR1, driven through productive contacts with NCOR2 in the inverse agonized state^17,18^.

Herein, we described our efforts to discover and characterize a covalent inverse agonist of PPARG suitable for clinical development in advanced UC.

## Results

RXRA induces an activation bias in PPARG that is enhanced by UC missense mutations and opposes inverse agonist efficacy

The previously described ability of RXRA^S427F^ to stabilize the active conformation of H12 in PPARG prompted us to explore the influence of RXRA^WT^ on PPARG in the context of TR-FRET assays modified from those described previously^4^. Briefly, we optimized conditions to ensure quantitative sensitivity to changes in the relative population of PPARG conformational states competent for binding co-factors (Supplementary Figure 1).

We initially probed the influence of RXRA^WT^ on the affinity of PPARG for a MED1 LxxLL peptide motif^4^ (CoA) and the NCOR1 ID2 CoRNR box^4,19^ (CoR). We found that addition of RXRA^WT^ resulted in a >10- fold enhancement of the affinity for MED1 while the affinity for NCOR1 was unchanged (Figure 1a and 1b) (RXRA^WT^:PPARG MED1 K_d_ 270 ± 4 nM vs >2,500 nM for PPARG alone; RXRA^WT^:PPARG NCOR1 K_d_ 340 ± 40 nM vs 390 ± 3 nM for PPARG alone). An increase in the basal affinity of the PPARG LBD for a CBP-derived CoA peptide in the presence of RXRA has been previously observed^20^, but to our knowledge, this is the first quantitative assessment of RXRA-enhanced binding of PPARG to a CoA motif. These results indicate that RXRA^WT^-PPARG heterodimers are equally poised for activation and repression in response to changes in the cellular environment (Figure 1c). This contrasts with conclusions drawn previously from studies of PPARG co-factor affinity in the absence of RXRA^5,21–23^. Further, we found that RXRA^S427F^ not only enhanced the affinity for CoA as seen previously^24^, but the greater plateau of the TR-FRET response indicates this mutation also bestows a greater fraction of PPARG to be competent for binding of CoA (Figure 1a). Intriguingly, we also found this mutation reduced the affinity for NCOR1 and that the relative reduction in CoR affinity is greater than the relative enhancement in CoA affinity (Figure 1a and 1b) (RXRA^WT^:PPARG NCOR1 K_d_ 340 ± 40 nM vs 1,200 ± 70 nM for RXRA^S427F^:PPARG), suggesting the primary biochemical role of this mutation may be to prevent CoR binding. Taken together, these results indicate that the conformational bias bestowed by RXRA^S427F^ drives the population of RXRA:PPARG heterodimers from being equally poised for CoA/CoR binding in the context of RXRA^WT^ heterodimers to having a significant activation bias that not only enhances co-activator recruitment but also significantly reduces co-repressor recruitment (Figure 1c).

**Figure 1.**
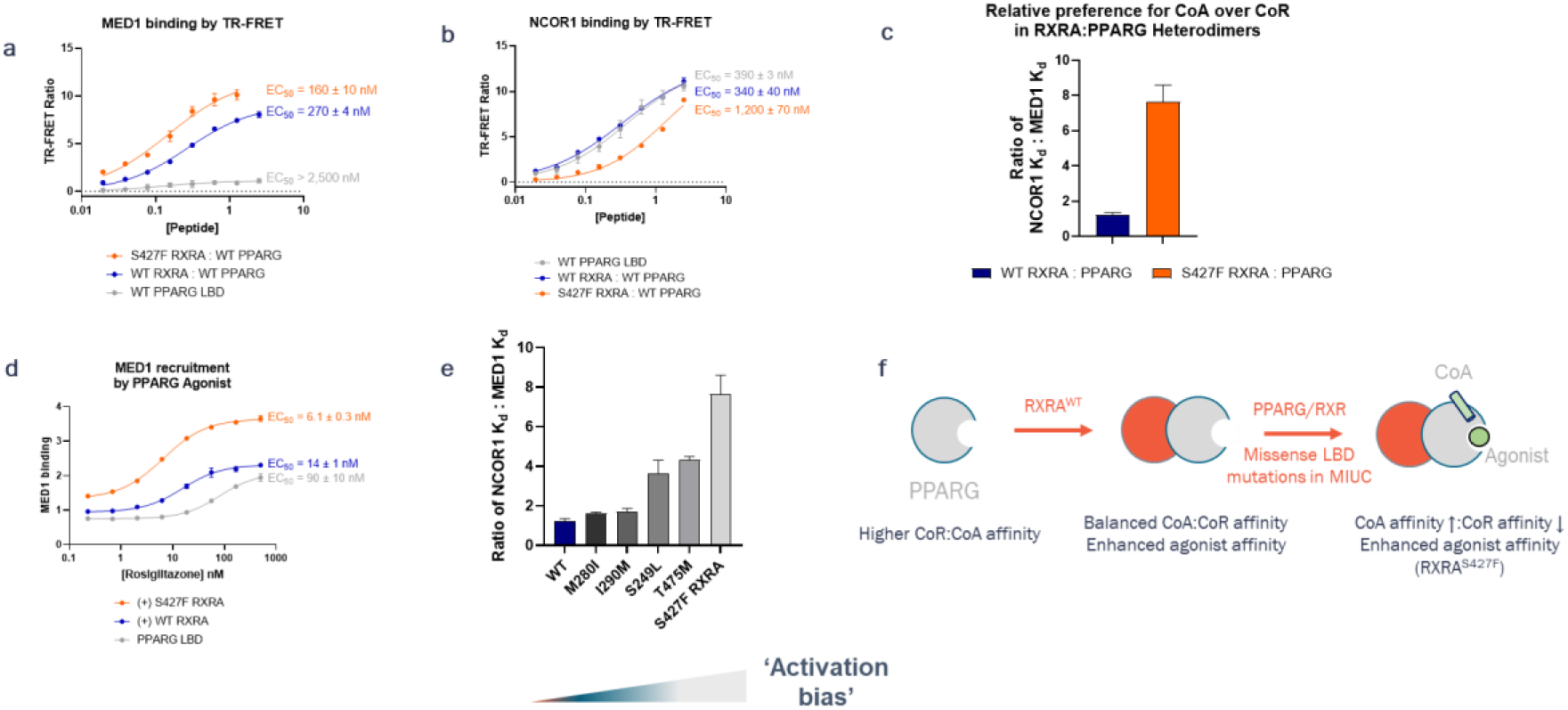
RXRA induces a conformationally activating bias in PPARG that is enhanced by MIUC mutations in PPARG and especially RXRA. (a) Binding affinity of a MED1 LxxLL Motif to PPARG LBD (gray), RXRA^WT^:PPARG^WT^ (blue) and RXRA^S427F^ (orange). Plots of representative curves from a single biological replicate done in technical duplicate are shown. Reported values are the average of two biological replicates ± the standard error of the mean (S.E.M.). Each biological replicate was performed in technical duplicate. (b) Binding affinity of NCOR1 ID2 Motif to PPARG LBD (gray), RXRA^WT^:PPARG^WT^ (blue) and RXRA^S427F^ (orange). Plots of representative curves from a single biological replicate done in technical duplicate are shown. Reported values are the average of two biological replicates ± S.E.M. Each biological replicate was performed in technical duplicate. Each biological replicate was performed in technical duplicate. (c) Relative affinities of RXRA^WT^:PPARG and RXRA^S427F^ heterodimers for a MED1 LxxLL motif and NCOR1 ID2, demonstrating an 8x conformational activation bias drive by RXRA^S427F^. Ratios were determined by averaging the ratio of the measured MED1 and NCOR1 K_d_ values ran side-by-side in two biological replicates. Each biological replicate was performed in technical duplicate. Error bars represent the S.E.M. between the biological replicates. (d) Rosiglitazone potency and maximal effect for CoA recruitment comparison across PPARG LBD, RXRA^WT^:PPARG and RXRA^S427F^:PPARG LBD heterodimers. Plots of representative curves from a single biological replicate done in technical duplicate are shown. Reported values are the average of two biological replicates ± S.E.M. Each biological replicate was performed in technical duplicate. (e) Comparison of ratio of CoA:CoR affinities across various MIUC mutant RXRA:PPARG LBD heterodimers. Ratios were determined by averaging the ratio of the measured MED1 and NCOR1 K_d_ values ran side-by-side in two biological replicates. Each biological replicate was performed in technical duplicate. Error bars represent the S.E.M. between the biological replicates. (f) Cartoon depiction summarizing biochemical findings on the influence of RXRA for driving PPARG to higher ligand sensitivity and balanced CoA/CoR affinity and the convergent function of various MIUC mutations to push this balance towards transcriptional activation through enhanced CoA relative to CoR affinity.

We also investigated the influence of RXRA^WT^ and RXRA^S427F^ on agonist-induced CoA binding utilizing rosiglitazone as a model ligand. We found that RXRA^WT^ enhanced the affinity of PPARG for rosiglitazone by approximately 5-fold and RXRA^S427F^ enhanced the affinity approximately 2-fold further than RXRA^WT^. Further, analogous to the greater plateau observed with titration of CoA, we observed RXRA^S427F^ enhancing the fraction of PPARG competent for binding CoA as evidenced by the E_max_ of the rosiglitazone dose-response curve (Figure 1d) (PPARG EC_50_ 90 ± 10 nM vs RXRA^WT^:PPARG EC_50_ 14 ± 1 nM vs RXRA^S427F^:PPARG EC_50_ 6.1 ± 0.3 nM). These findings indicate that a previously unappreciated role of the RXRA^S427F^ mutation is also to enhance the response of PPARG to its endogenous ligand(s) in UC.

Across a panel of muscle invasive UC (MIUC) missense mutations in PPARG and RXRA^12^, we found a unifying theme of biasing of the conformational landscape towards activation through alterations of the relative affinities for co-activator and co-repressor (Figure 1e and Supplementary Figure 2). PPARG^S249L^ proved the most unique by almost exclusively driving an activation bias through repulsion of CoR binding (Supplementary Figure 2). Taken with the influence of RXRA^S427F^ on CoR binding, this mutation may be hinting at the importance of disfavoring CoR binding in PPARG-activated MIUC. A summary of our biochemical learnings on RXRA/PPARG missense mutations is depicted in Figure 1f. Finally and likely not coincidentally, the most powerful mutations to drive an activation bias in PPARG are also the most clinically prevalent^12^, namely, RXRA^S427F^ and PPARG^T475M^, with RXRA^S427F^ driving the most significant activation bias biochemically while also being the most clinically recurrent. We therefore focused our additional studies on this mutation.

Optimization of repressive biasing in PPARG to counter RXRA^S427F^-induced activation bias

Informed by our mutational work on MIUC missense mutations, especially RXRA^S427F^ (Figure 1), we hypothesized that modulators that induce an enhanced repressive bias in PPARG:RXRA^WT^ heterodimers could counter the activation bias bestowed by RXRA^S427F^. Owing to the unique conformational state induced by T0070907^16^, we chose to optimize covalent modifiers of PPARG to counter the MIUC activation bias observed biochemically (Figure 1). Based on the observation that T0070907 and GW9662^25^ have a similar potency in disrupting agonist-induced CoA recruitment but differential abilities to both repel basal CoA recruitment and induce CoR recruitment (Supplementary Figure 3), we surmised that the conformation of covalent inverse agonists that results in covalent capture of C313 is distinct from the conformational state that exhibits high affinity for CoR. This was independently corroborated by crystallographic studies utilizing C313A and a T0070907 analog^17^. Therefore, to pharmacologically counter the MIUC activation bias, our primary medicinal chemistry aim was to optimize stabilization of the unique, high affinity conformational state for CoR (E_max_). Biochemical potency (EC_50_) was a secondary focus.

In contrast to previous efforts^18^, we elected not to target *de novo* interactions with any specific CoR peptide but instead to focus on reinforcement of the PPARG conformational state, which should theoretically bestow high affinity for all CoR’s. This strategy could avoid challenges in maintaining efficacy due to alterations in the balance of expression of diverse CoR’s. Structure-based drug design of the T0070907-PPARG adduct (PDB: 6ONI) in the context of the high affinity state for CoR led us to hypothesize the following: 1) conformational constraint of the amide moiety of T0070907 would pre-organize the adduct to the appropriate conformation to enhance CoR recruitment and 2) enhancement of the N-H acidity would strengthen the H-bond with the C-terminal carboxylate of PPARG Y505, stabilizing the insertion of H12 into the ligand-binding pocket, thereby enhancing CoR recruitment. Additionally, we sought to replace the nitro group with the cyano S_N_Ar activating group to reduce general reactivity and to improve metabolic stability. After several iterations, this strategy led us to the identification of FTX-6746 (Figure 2a).

**Figure 2.**
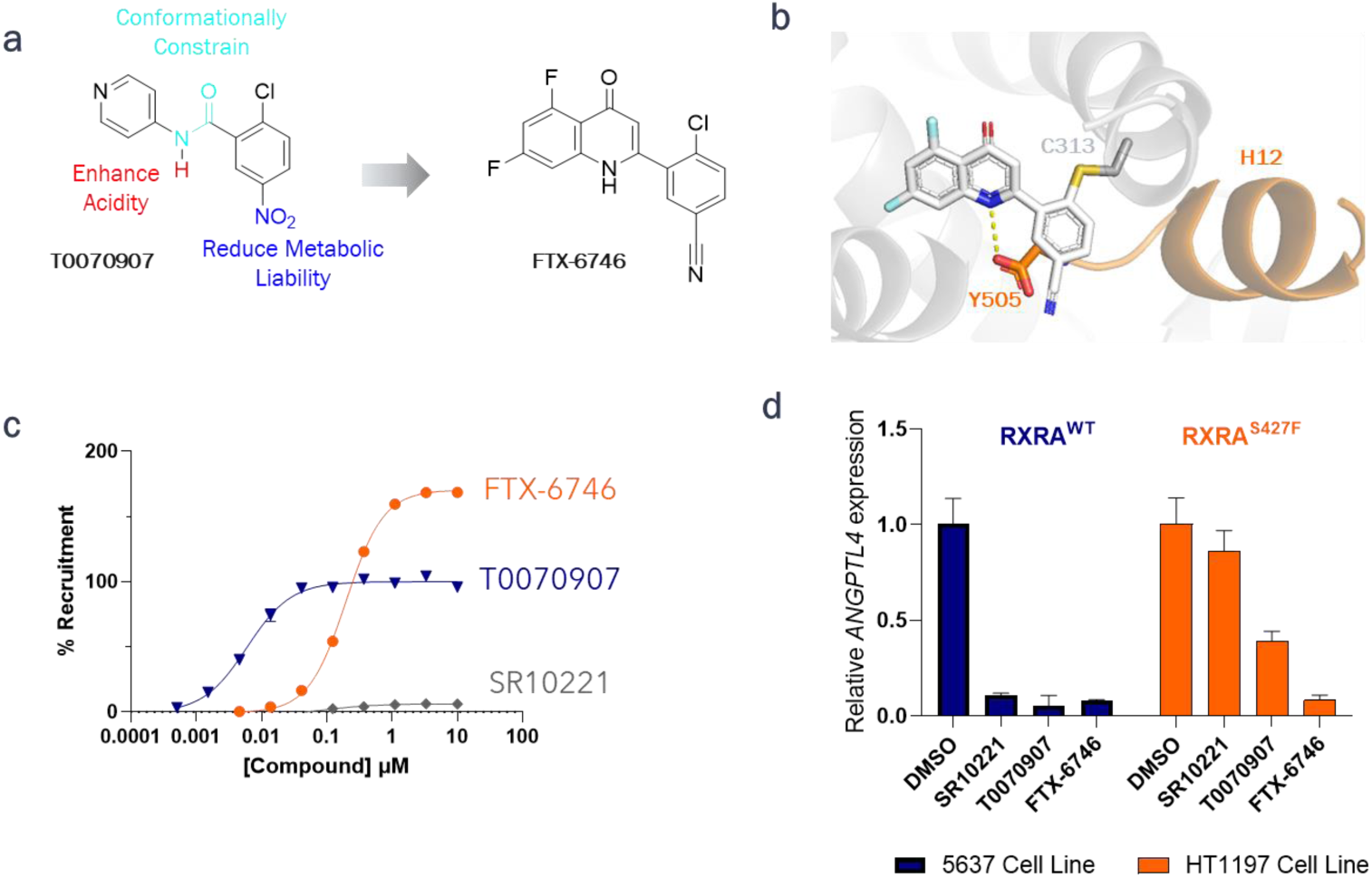
Optimization of repressive biasing in PPARG to counter RXRA-induced conformational biasing. (a) Overview of medicinal chemistry strategy for covalent PPARG inverse agonist optimization resulting in FTX-6746. (b) Crystal structure of FTX-6746 covalent adduct with PPARG LBD. The PPARG LBD (gray) adopts a similar conformation to that observed with T0070907 (PDB 6ONI) with H12 (orange) inserted into the ligand binding pocket with the C-terminal carboxylate forming a key H-bond with the quinolone nitrogen of the FTX-6746 adduct (yellow dashes). (c) Dose-dependent recruitment of NCOR1 ID2 to RXRA^S427F^:PPARG heterodimers, demonstrating the superiority of FTX-6746 for countering the activation bias driven by this mutation over T0070907 and SR10221. Representative curves from ≥ 3 biological replicates are shown. Data points indicate the average of technical duplicates from a representative biological replicate. The 95% confidence interval for EC_50_ values of SR10221, T0070907 and FTX-6746, respectively, were determined to be: 109 nM ± 30 nM, 60.5 ± nM, 197.5 ± 8.5 nM. The 95% confidence interval for the E_max_ values relative to T0070907 for SR10221 and FTX- 6746, respectively, were determined to be: 6.2 ± 0.6% and 170.5 ± 2%. Rosiglitazone was not present for these experiments. (d) Comparison of SR10221, T007097 and FTX-6746 ability to suppress PPARG target gene *ANGPTL4* in RXRA^WT^ (blue, 5637 cell line) and RXRA^S427F^ (orange, HT1197 cell line) contexts. Data indicate the activation bias observed biochemically reduces T0070907 and SR10221 efficacy and the superior repressive biasing of FTX-6746 enables it to achieve similar repressive effects in both contexts. Data is representative of two separate biological experiments.

The structure of the covalent adduct of FTX-6746 with PPARG and NCOR1 was solved using x-ray crystallography and as expected, we found that the adduct reinforced a similar conformation as that seen previously with T0070907 (Figure 2b). To study the overall conformational biasing in solution, we characterized the PPARG-FTX-6746 adduct using both HDX and alterations in CoA binding by surface plasmon resonance (SPR) analysis (Supplementary Figure 4). These studies confirmed the conformational landscape of PPARG was changed without relying on CoR-specific interactions. In the context of HDX, enhanced shielding throughout H12 was observed relative to T0070907, indicating a change in the conformational landscape of PPARG with the data supporting increased residence time and/or increased sampling of the H12 insertion into the PPARG ligand-binding pocket. By SPR, FTX- 6746 adduct formation resulted in a decreased fraction of PPARG capable of binding CoA (82% vs 34%, apo vs. FTX-6746-modified, respectively) also supporting a change in the conformational balance of PPARG covalently modified by FTX-6746.

To study the functional consequences of enhanced repressive biasing of FTX-6746, we utilized both T0070907 and SR10221, to compare to a prototypical covalent and noncovalent inverse agonist, respectively. Consistent with previous studies, we found a dramatic difference in the E_max_ of recruitment between T0070907 and SR10221^11^. Strikingly, we observed that the E_max_ for NCOR1 recruitment to PPARG in the presence of RXRA^S427F^ for both compounds was significantly decreased relative to RXRA^WT^. Further, we found a similar trend for the ability of these compounds to repel MED1 binding to RXRA^WT^ and RXRA^S427F^ heterodimers, further increasing our confidence in the significance of this observation (Supplementary Figure 5). Importantly, we observed that FTX-6746 exhibited superior conformational biasing of PPARG as determined by the E_max_ of the dose response curve for NCOR1 recruitment to both RXRA^WT^ and RXRA^S427F^ heterodimers (Supplementary Figure 6 and Figure 2c respectively) (E_max_ differential of FTX-6746 relative to T0070907 was 176% ± 6% and 170.5% ± 2% for RXRA^WT^- and RXRA^S427^–complexes with PPARG, respectively).

To validate the biological significance of our biochemical findings, we compared the ability of these molecules to suppress a PPARG target gene in an RXRA^WT^ UC cell line (5637) and an RXRA^S427F^ UC cell line (HT1197). Similar to the observation that RXRA^S427F^ reduced the ability of T0070907 and SR10221 to recruit co-repressors, it also reduced the efficacy of both compounds to suppress *ANGPTL4* transcript levels in RXRA^S427F^ cells relative to RXRA^WT^ cells. Further, the enhanced conformational biasing seen with FTX-6746 resulted in superior PPARG target gene suppression in both RXRA^WT^ and RXRA^S427F^ cellular contexts compared to T0070907 and SR10221 (Figure 2d). This enhanced suppression is decoupled from the biochemical potency but is correlated with the E_max_ for biochemical CoR recruitment, supporting our mechanistic model separating binding, reactivity, and conformational biasing, while underscoring the centrality of conformational biasing for inverse agonist optimization. Finally, the biochemical-biological correlation was restricted to the E_max_ for NCOR1 recruitment and not MED1 blockade, at which SR10221 was biochemically superior but nonetheless biologically inferior, supporting our earlier observations that MIUC mutations may primarily repel CoR binding over CoA recruitment (Figure 1a, 1b, 2c, 2d and Supplementary Figures 6)

### Utilizing mechanism to enable *in vivo* optimization of PPARG covalent inverse agonists to discover FX- 909

During our optimization, we observed that both FTX-6746 and T0070907 displayed low intrinsic reactivity toward nucleophilic aromatic substitution (S_N_Ar) with glutathione (GSH) under the conditions tested, even out to 24 hours. In contrast, both compounds show rapid reactivity with C313 in PPARG. (Supplementary Figure 7a). Further hinting at the atypical nature of S_N_Ar capture of C313, we observed a reverse trend in halogen reactivity on the quinolone scaffold of FTX-6746 compared to a standard nucleophilic aromatic substitution (S_N_Ar) reaction^26^ (-F>-Cl>-Br; Supplementary Figure 7b), similar to previous findings^18^. To probe the mechanistic basis for these observations, we evaluated the temperature-dependence of the reaction rate constant 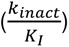 of FTX-6746 and T0070907 with PPARG C313. Strikingly, we found that we could not reasonably fit our experimental data to the Eyring equation, suggesting a reaction mechanism with multiple transition states (Supplementary Figure 7c and 7d).

The totality of these observations in Supplementary Figure 7a-d indicated to us that the S_N_Ar reaction of these compounds with PPARG-C313 proceeds via a stepwise process, thus explaining the existence of multiple transition states along the reaction coordinate. This would suggest that the reversible affinity (*K_I_*) of these modifiers comprises not only noncovalent interactions as is typically assumed for this value but also includes the stability of the tetrahedral intermediate (Meisenheimer Complex) accessed during stepwise S_N_Ar reaction mechanisms (Figure 3 and Supplementary Figure 7e). Our interpretation of these mechanistic studies suggested to us that non-halogen based leaving groups could be utilized for the optimization of the S_N_Ar reaction while retaining the identical covalent adduct as FTX-6746 for equivalent repressive conformational biasing. This approach led to FX-909, which contains a more strongly electron-withdrawing methylsulfone leaving group that more stably exists in the Meisenheimer Complex with PPARG, reflected in the lower *K_I_* of FX-909 (*K_I_* = 2.7 ± 0.7 µM) compared to FTX-6746 (*K_I_* = 12 ± 6 µM) while maintaining a similar *k_inact_* (Figure 3b) (0.33 ± 0.09 min^- 1^ vs 0.332 ± 0.004 min^-1^; FTX-6746 vs FX-909). Crucially, the overall modest improvement in the biochemical potency of FX-909 for PPARG was accompanied by improved *in vivo* properties of the molecule relative to FTX-6746 (Supplementary Table 1).

**Figure 3.**
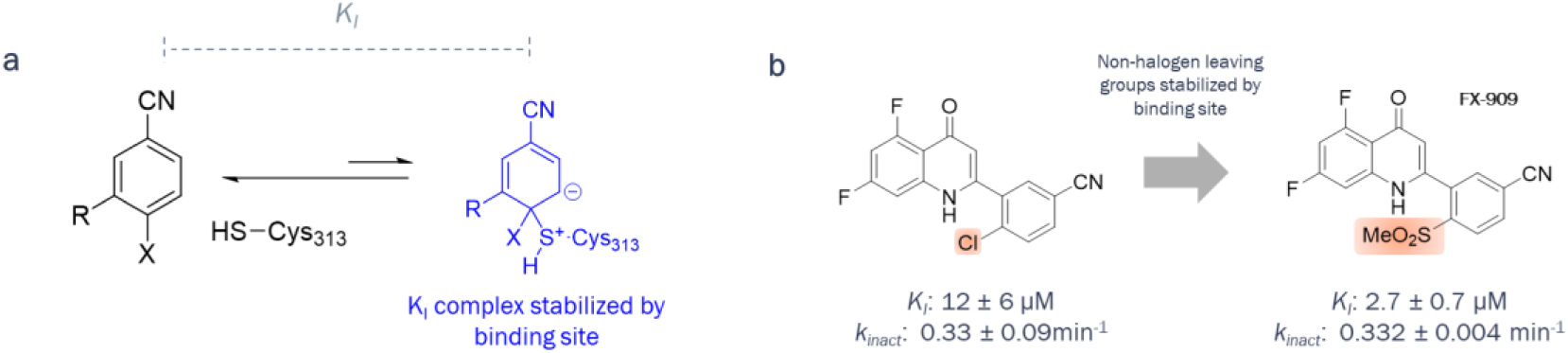
Mechanistic understanding of S_N_Ar capture of C313 to provoke leaving group SAR exploration resulting in identical adduct formation as FTX-6746 with superior PK properties. (a) Meisenheimer Complex is key intermediate species contributing to K_I_ in S_N_Ar capture of C313 as indicated by halogen leaving group SAR and from temperature-dependent reaction rate studies (see Supplementary Figure 7). (b) Methylsulfone leaving group leads enhancement of K_I_ and not k_inact_, attributed to enhanced formation and stabilization of the Meisenheimer Complex in the binding site. Reported *KI* and *kinact* values for FTX-6746 are the average ± S.D. of 3 biological replicates each performed as technical duplicates. Values for FX-909 are the average ± S.E.M. of 2 biological replicates, each performed as technical duplicates.

### Molecular characterization of FX-909

The context-dependent electrophilicity observed of the 4-(methylsulfonyl)benzonitrile warhead with PPARG suggested FX-909 would exhibit exquisite selectivity across the proteome. To test this, we first evaluated the selectivity of FX-909 for PPARG against PPARA and PPARD, with >2,000-fold selectivity observed for PPARG (Supplementary Figure 8). To assess selectivity proteome-wide, mass spectrometry (MS) analysis was used to assess FX-909 cysteine reactivity using three UC cell lines (UMUC9, HT1197 and 5637). FX-909 displayed specific engagement on PPARG C313 following treatment in all three cell lines with minimal evidence of off-target labeling (Figure 4a). Similar proteome-wide selectivity was observed with FTX-6746 (Supplementary Figure 9)

**Figure 4.**
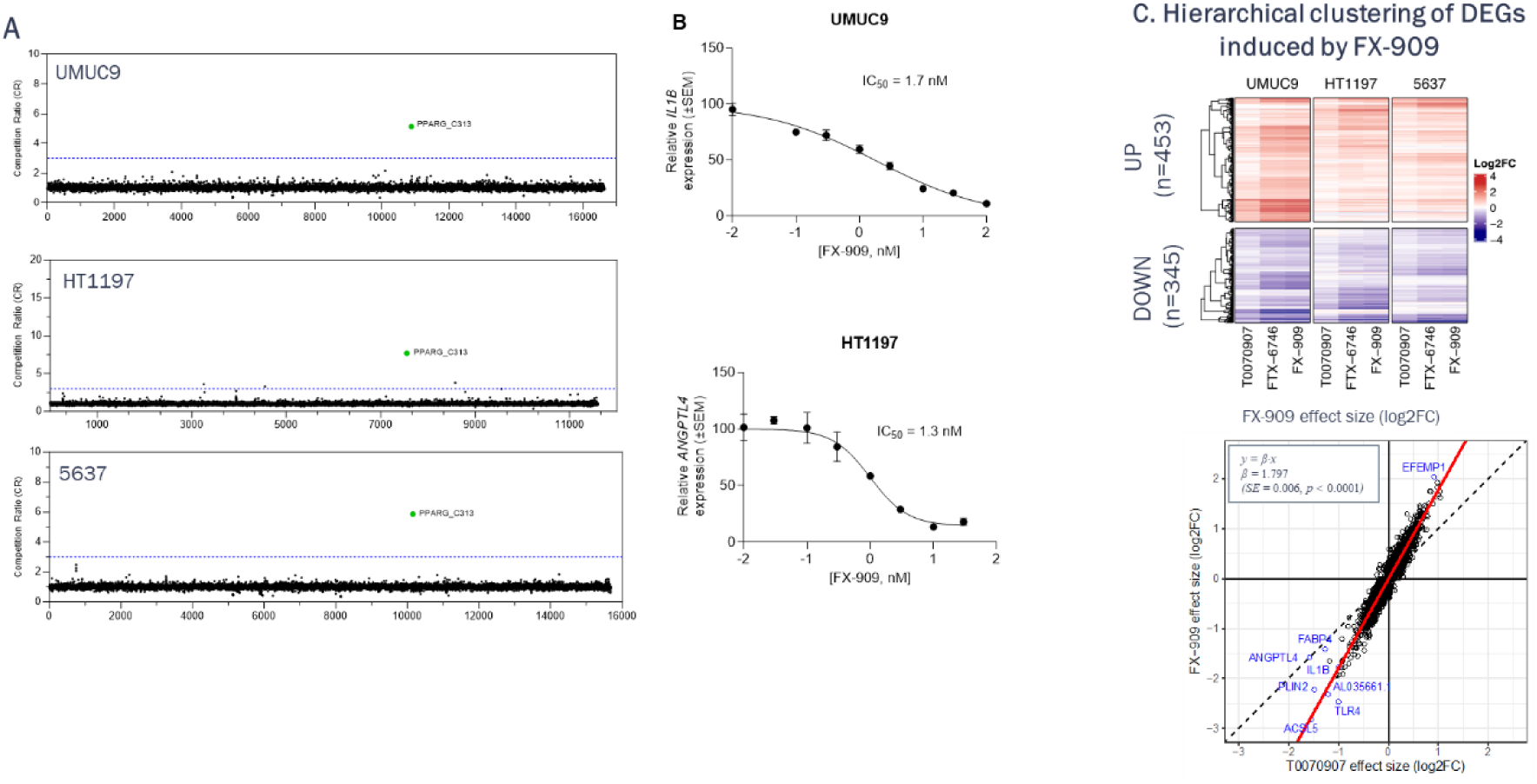
(A) Selective engagement of FX-909 across urothelial carcinoma cell lines. Global cysteine profiling and engagement of FX-909 in UMUC9, HT1197 and 5637 cell lines following 30 min treatment. PPARG C313 was the only significantly engaged target (orange). Competition ratio (CR) was calculated dividing the control channel (DMSO) by the electrophile treated channel. The blue dotted line indicates a CR of 3. (B) Cellular potency of FX-909. (top) Dose-dependent suppression of *IL1B* expression by FX-909 in the urothelial carcinoma line UMUC9. Relative *IL1B* expression and IC_50_ value are indicated. (bottom) Dose-dependent suppression of *ANGPTL4* expression by FX-909 in the urothelial carcinoma line HT1197. Relative *ANGPTL4* expression and IC_50_ value are indicated. (C) Transcriptional changes induced by PPARG inverse agonists in UMUC9, HT1197 and 5637 cell lines following 24 hr treatment. (top) Heatmap showing the effect of T0070907, FTX-6746 and FX-909 on the genes that are differentially up- or down-regulated by FX-909 (see Supplementary Figure 10). (bottom) Comparison of gene expression changes induced by FX-909 vs T0070907. Each dot corresponds to a gene that showed statistically significant change in response to T0070907, FTX- 6746 or FX-909 (n=6,759 genes; FDR < 0.05) as assessed from a mixed effect model where cell lines are treated as a random effect. Red line corresponds to a linear regression model with a slope of β=1.8, indicating that on average FX-909 induces 3.5=fold (2^1.8) times greater fold changes in gene expression. Dashed black line shows where data would fall if FX-909 and T0070907 induced effects of the same magnitude. The lack of gene expression changes in the bottom right and top left quadrants of the plot indicates that FX-909 does not regulate different genes from T0070907.

To assess the ability of FX-909 to modulate PPARG function in cells, PPARG target gene expression was analyzed in UC cell lines after treatment with FX-909. FX-909 suppressed *IL1B* mRNA by 50% (inhibitory concentration at 50%, IC_50_) at a concentration of 1.7 nM in the UMUC9 cell line and suppressed *ANGPTL4* mRNA with an IC_50_ of 1.3 nM in the HT1197 cell line (Figure 4b). Different PPARG target genes were chosen for different cell lines based on observation of robust, dose-dependent effects.

To evaluate the global transcriptomic effects, RNA sequencing of UMUC9, HT1197 and 5637 cell lines was also performed after treatment with either FX-909, FTX-6746 or T0070907. Principal component analysis (PCA) of the treatment-induced effects revealed that, while different genes were regulated differently in different cell lines, across all three cell lines, the magnitude of PPARG-regulated gene expression changes was higher when treated with either FX-909 or FTX-6746 than when treated with T0070907 (Supplementary Figure 10). To obtain a consensus FX-909-induced gene expression profile, transcriptomics data across all cell lines were analyzed using a linear mixed effects model (Figure 4c; Supplementary Figure 10). When comparing the consensus FX-909 to the consensus T0070907 gene expression changes, FX-909 overall showed 1.8-times greater effect than T0070907 (Figure 4c). Cut&Run analysis performed on UMUC9 cells treated with DMSO or FTX-6746 showed reduced binding of PPARG and RXRA at PPARG target gene loci, including *ANGPTL4* and *FABP4* (Supplementary Figure 11).

GSEA pathway analysis shows that FX-909 results in downregulation of cell proliferation-related pathways (Supplemental Figure 12a), consistent with its anti-proliferative effects (see below). FX-909 also results in downregulation of adipogenesis and fatty acid metabolism pathways (Supplementary Figure 12b) as well as genes upregulated by the PPARG agonist, Rosiglitazone (Supplementary Figure 12c), confirming FX-909’s mechanism of action via PPARG inverse agonism. Among upregulated gene sets are components of the complement system, EMT genes and genes involved in extracellular matrix organization (Supplementary Figures 12a and 12b).

### *In vitro* efficacy of PPARG inverse agonists in UC models

Phenotypic analysis of PPARG inverse agonists was performed in a panel of 14 UC cell lines. Using clonogenic growth assays, FX-909, FTX-6746 and T0070907 were treated with a range of concentrations to generate a GI_50_ value. GI_50_ values for FX-909 ranged from 6 nM to >500 nM while T0070907 treatment did not result in 50% growth inhibition in any cell lines except UBLC1, where the GI_50_ was modest relative to that of FX-909 (∼250 nM vs 6 nM, respectively) (Figure 5a, Supplementary Figure 13). Cells that are characterized as PPARG pathway activated – mutated or amplified for PPARG, RXRA mutated, or high expression of PPARG pathway gene signature^12^ showed preferential sensitivity to FX-909 and FTX-6746 over T0070907.

**Figure 5.**
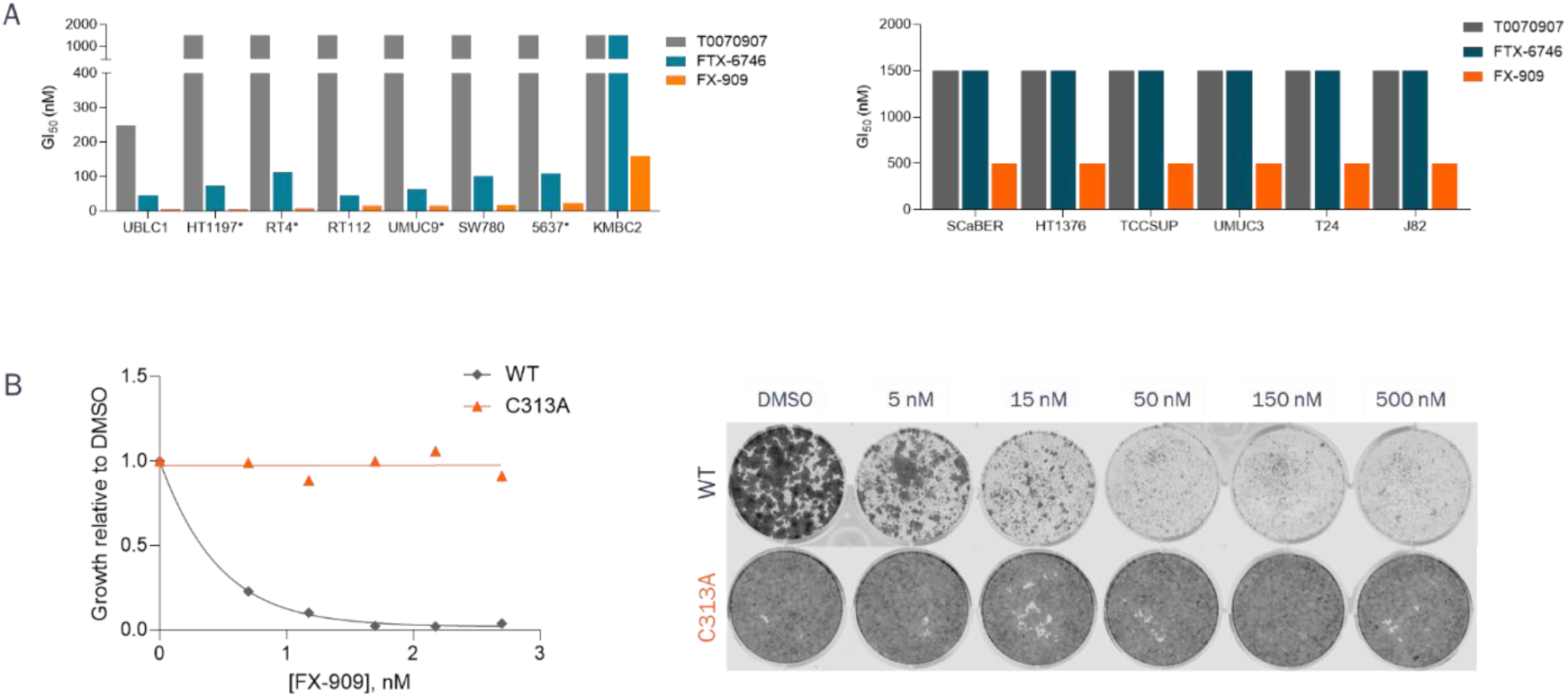
(A) Phenotypic potency of PPARG inverse agonists. GI_50_ values of UC cell lines treated with T0070907, FTX-6746 or FX-909 for 11-14 days. GI_50_ values were calculated relative to the DMSO treated cells using a crystal violet stain quantitation. GI_50_ values greater than the maximum dose tested - 500 nM (FX-909) or 1500 nM (T0070907 and FTX-6746) - were graphed as 500 nM and 1500 nM, respectively. * = PPARG/RXRA mutant or PPARG amplified lines. Data are a representative of multiple individual experiments performed. (B) Phenotypic potency of FX-909 in WT or C313A isogenic HT1197 cell lines. Treatment of parental and C313A HT1997 cells lines for 14 days with FX-909 resulted in potent growth inhibition only in parental line, growth curve on left and images of clonogenic assay on right.

To determine the specificity of FX-909-mediated phenotypic growth effects, isogenic HT1197 cells were generated by knock-in of a C313A mutation in PPARG, which prevents the covalent modification of PPARG by FX-909. While PPARG^C313A^ knock-in cells grow more slowly than parental lines, treatment of the parental or knock-in cell line with a dose titration of FX-909 illustrated the specificity of the effects as C313A mutant cells were completely resistant to FX-909 growth effects (Figures 5b). Of note, the anti-proliferative effects of FX-909 occur at concentrations similar to those required for the suppression of PPARG target gene suppression (Figure 4b).

### *In vivo* studies of PPARG inverse agonists in UC models

*In vivo* studies to assess the effects of PPARG inverse agonists on tumor growth were performed with two UC xenograft models, UMUC9 and HT1197. Mice harboring xenograft tumors were dosed with vehicle, FTX-6746 or FX-909. UMUC9 xenografts were sensitive to FTX-6746, with similar tumor regression observed after 21 days of treatment with 30 or 60 mg/kg given orally (PO) twice a day (BID). FX-909 oral administration at 3, 10 or 30 mg/kg BID also resulted in tumor regressions that were greater than those observed with FTX-6746, even in the FX-909 3 mg/kg dose group (108% vs 119% TGI) (Figure 6a). After 21 days of dosing, a subset of animals was observed for tumor regrowth for an additional 45 days in the absence of treatment. While the FTX-6746-treated animals showed tumor regrowth upon drug cessation, the FX-909 animals showed negligible tumor regrowth during the observation period. HT1197 xenografts were also sensitive to FTX-6746 (60 mg/kg BID) and FX-909 (1, 3, 10 mg/kg BID) administration, with tumor regression observed at all FX-909 doses. Similar tumor regressions were observed with FTX-6746 at 60 mg/kg BID and FX-909 at 1 mg/kg BID dose (130% TGI vs 121% TGI, respectively) while higher doses of FX-909 resulted in deeper regressions (151% at 10 mg/kg BID and 166% at 30 mg/kg BID) (Figure 6b). All drug doses in both models were tolerated as measured by body weight, with no animals losing >10% body weight during the dosing period (data not shown).

**Figure 6.**
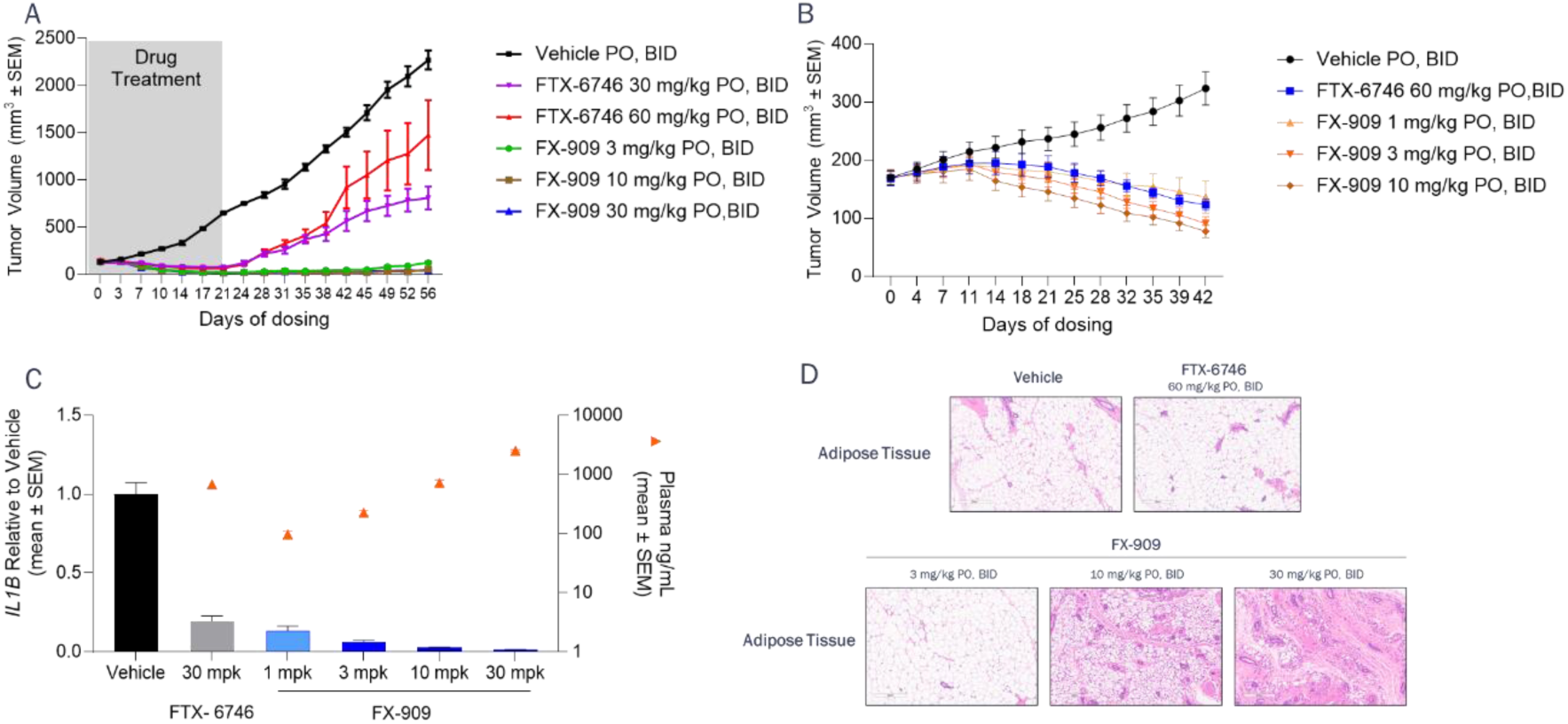
– (A) UMUC9 xenograft efficacy study shows FX-909 elicits durable tumor regression at all doses, while FTX-6746 elicits tumor regression that is not durable upon drug cessation. (B) HT1197 xenograft efficacy study shows tumor regression for both FX-909 and FTX-6746. (C) PK/PD relationship for FTX-6746 and FX-909. (D) H&E staining of adipose tissue from mice in study shown in (A).

Pharmacokinetic (PK)/pharmacodynamic (PD) analysis was also performed for PPARG inverse agonists in the UMUC9 xenograft model. Mice were dosed orally with three BID doses of vehicle, FTX- 6746 (30 mg/kg) or FX-909 (1, 3, 10 or 30 mg/kg) prior to harvest of plasma and tumor for analysis. Tumor PD was assessed using qPCR to quantitate changes in PPARG target gene *IL1B* and PK analysis was performed on plasma to quantitate exposure (Figure 6c). FTX-6746 treatment resulted in robust suppression of *IL1B* in the tumor samples, with 80% reduction in *IL1B* after administration of 30 mg/kg of FTX-6746. FX-909 resulted in 86% suppression of *IL1B* at the lowest dose tested (1 mg/kg), with increasing suppression observed with higher doses and near complete suppression at 30 mg/kg (99%). PK analysis showed similar exposure for 30 mg/kg FTX-6746 and 10 mg/kg FX-909. FX-909 exposures increased in a dose proportional manner.

As PPARG is known to be a key regulator of adipocyte biology, inguinal white adipose tissue was collected from animals in the UMUC9 efficacy study at the end of dosing on day 21. Formalin fixed paraffin embedded (FFPE) tissue samples were sectioned and stained with hematoxylin and eosin (H&E) (Figure 6d). Adipose tissue from 60 mg/kg BID FTX-6746 and 3 mg/kg BID FX-909 treated animals exhibited similar morphology to the vehicle control animal tissue, while higher doses of FX- 909 (10 and 30 mg/kg BID) showed alterations in the morphology of the tissues with obvious loss of normal adipocytes, indicating on-target effects of PPARG inhibition, similar to those observed with knockout of PPARG^27,28^.

## Discussion

Our studies in pursuit of PPARG covalent inverse agonists for the treatment of patients with MIUC revealed a number of fundamental mechanistic relationships.

First, dimerization of PPARG with RXRA^WT^ is sufficient to enhance the affinity of PPARG for rosiglitazone and therefore, likely all agonist-type ligands. RXRA^WT^ also balances the conformational landscape of PPARG to be poised for activation or repression. This balancing, combined with the affinity enhancement for agonist by RXRA^WT^ provides a mechanistic basis for the oncogenic role of PPARG amplification and overexpression in MIUC whereby elevated levels of PPARG compete for the limited pool of RXRA present, sensitizing the cells to PPARG endogenous ligands to drive MIUC growth.

Second, the RXRA^S427F^ hotspot mutation not only enhances the affinity of PPARG for CoA peptides as previously described^24^, but also enhances the cooperativity between PPARG and RXRA to further sensitize these heterodimers to agonists. This mutation also reduces the affinity of the heterodimer for a CoR peptide, creating an activation bias in the heterodimer conformational landscape to drive the PPARG transcriptional program. This bias can be seen with all PPARG LBD mutations tested. Taken together, our characterization of both RXRA^WT^:PPARG and RXRA^S427F^:PPARG heterodimers present a unifying biochemical mechanistic framework by which various genetic and epigenetic events activate the PPARG gene program to drive MIUC.

Third, our understanding of the RXRA:PPARG conformational landscape in MIUC allowed us to rationally tune the biological activity of PPARG covalent inverse agonists to reverse the activation bias present in MIUC to a repressive bias to suppress the PPARG transcriptional program in cells.

Fourth, our dissection of the mechanistic pathway of S_N_Ar covalent modifiers for PPARG elucidated an atypical mechanism in which reversible binding (*K_I_*), reactivity (*k_inact_*) and subsequent conformational biasing are each independently tunable to optimize biological activity. Further, the context-dependence of the S_N_Ar reaction allows for exquisite selectivity from this 4-(methylsulfonyl)benzonitrile warhead in biological contexts and the stepwise nature of this mechanism provides additional tuning parameters for *in vitro* and *in vivo* optimization.

This series of mechanistic insights culminated in FX-909, a highly selective covalent inverse agonist of PPARG. FX-909 more stably enforces a repressive conformation in PPARG that enhances its affinity for co-repressors and decreases its affinity for co-activators, thereby driving both potent and deep suppression of PPARG target genes to inhibit PPARG-mediated MIUC growth both *in vitro* and *in vivo*. FX-909 potency, *in vivo* properties, and depth of target gene suppression result in durable regression in both RXRA^S427F^ and PPARG-amplified MIUC xenograft models.

FX-909 represents a promising therapeutic agent for the treatment of PPARG-activated urothelial cancers. It is currently being assessed in a phase 1 clinical trial (NCT05929235).

## Methods

Commercial Compound Sourcing – T0070907, GW9662 were purchased from Cayman Chemical and SR10221 was SR10221 was synthesized as described previously^29^

### Recombinant protein expression and structure determination

#### Expression and purification of human PPARG for assay

The wild type and mutant (S249L, M280I, I290M and T475M) DNA of human PPARG LBD domain (residues 231 to 505) was cloned into pET21b expression vector with an N-terminal TEV-cleavable His6 tag. The plasmids were transformed into *Escherichia coli* BL21(DE3) gold strain (Novagen) for expression. Cells were grown at 37°C to an optical density of 0.6 to 0.8, and expression was induced by addition of 0.1 mM IPTG and overnight incubation at 16°C. Cells were collected by centrifugation and resuspended in lysis buffer (50mM Tris pH 8.0, 500mM NaCl, 5% glycerol). Cells were disrupted by sonication and lysate was prepared by centrifugation for 1h at 13,500rpm. After centrifugation, the lysate was isolated by Ni-NTA (Qiagen). PPARG protein was eluted using lysis buffer plus 250 mM imidazole, the eluted protein was concentrated and diluted by 10-fold buffer A (50mM Tris pH 8.0, 5% glycerol). The protein was purified by ion-exchange chromatography using a HiTrapTM Q-HP 5 mL column (Cytiva) and washed using a gradient of 0∼50% buffer B (50mM Tris pH 8.0, 1M NaCl 5% glycerol) over 20 column volumes. The fractions of interest were pooled together and purified by size exclusion chromatography using a Superdex 75 10/300 GL column (Cytiva) equilibrated with size-exclusion buffer (20mM Tris pH 8.0, 150mM NaCl, 10% glycerol, 2mM DTT). Pure protein fractions were pooled together and stored at -80°C.

#### Expression and purification of human PPARG for crystallization

The wild type DNA of Human PPARG LBD domain (residues 231 to 405) was clone into pET21b expression vector with an N-terminal TEV-cleavable His6 tag. The plasmids were transformed into *Escherichia coli* BL21(DE3) gold strain (Novagen) for expression. Cells were grown at 37°C to an optical density of 0.6 to 0.8, and expression was induced by addition of 0.1 mM IPTG and overnight incubation at 16°C. Cells were collected by centrifugation and resuspended in lysis buffer (50mM Tris pH 8.0, 500mM NaCl, 5% glycerol). Cells were disrupted by sonication and lysate was prepared by centrifugation for 1h at 13,500 rpm, 4°C. After centrifugation, the lysate was isolated by Ni-NTA (Qiagen). PPARG protein was eluted using lysis buffer plus 250mM imidazole, then dialyzed overnight against lysis buffer in the presence of HIS-TEV protease. Cleaved protein was loaded onto the Ni-NTA column again to remove HIS-TEV. The protein was purified by ion-exchange chromatography using a HiTrapTM Q-HP 5 mL column (Cytiva). The final protein was purified by size exclusion chromatography using a Hiload 10/600 superdex 75 (Cytiva) equilibrated with size-exclusion buffer (20mM KH2PO4- K_2_HPO_4_, pH 7.4, 50mM NaCl, 5mM TCEP, 0.5mM EDTA). Pure protein fractions were pooled together and concentrated to 15-20mg/ml and stored at -80°C.

Wild type PPARG (residues 231 to 505) was incubated with FTX-6746 for 36h at 1:5 protein/ligand molar ratio. NCOR1 (DPASNLGLEDIIRKALMGSFDDK) was then added, and the mixture was incubated for 6h at a 1:5 protein/peptide molar ratio and concentrated to 10 mg/mL for crystal wide screening and optimization. Crystals were grown by hanging drop vapor diffusion after 4-5 days under 0.1 M MES pH 6.5, 0.1M ammonium sulfate and 24 % w/v PEG 8000 at 18°C. The crystals were cryo- protected in mother liquor containing 15% glycerol and flash-frozen in liquid nitrogen on nylon loops.

#### Data collection and structure determination

The diffraction data was collected at Advanced Photon Source (APS 21-ID-D) and processed using XDS. The initial molecular replacement was performed by Phaser MR (PDB code: 6ONI) as the template. Structure refinement was carried out using Refmac5 combined with several rounds of COOT manual fitting. Figures were prepared with PyMOL (The PyMOL Molecular Graphics System). Complete data collection and refinement statistics are shown in Supplementary Table 2.

#### Expression and purification of human RXRA

The wild type and mutant (S427F) DNA sequences encoding human RXRA LBD domain (residues 228 to 462) with N-terminal FLAG and TEV tag were cloned into pET21b vector. The plasmids were transformed into *Escherichia coli* BL21(DE3) gold strain (Novagen) for expression. Cells were grown at 37°C to an optical density of 0.6 to 0.8, and expression was induced by addition of 0.1 mM IPTG and overnight incubation at 16°C. Cells were collected by centrifugation and resuspended in lysis buffer (20mM Tris pH 7.5, 500mM NaCl). Cells were disrupted by sonication and lysate was prepared by centrifugation for 1h at 13,500 rpm, the lysate was loaded on FLAG Resin (Sigma). Protein eluted with 0.2 mg/ml FLAG peptide was concentrated for size exclusion chromatography using a Superdex 75 10/300 GL column (Cytiva) and SEC buffer (20mM Tris pH 7.5, 500mM NaCl). Fractions containing pure protein were pooled together and stored at -80°C.

### TR-FRET assays

HTRF MAb Anti-6HIS Tb cryptate gold and HTRF streptavidin-d2 were obtained from Cisbio, as well as HTRF PPI terbium detection buffer. Co-repressor and co-activator peptide sequences used were biotinylated NCOR1 ID2 peptide (Biotin-GHSFADPASNLGLEDIIRKALMG-amide) and biotinylated MED1 LxxLL peptide (Biotin-VSSMAGNTKNHPMLMNLLKDNPAQ-amide), respectively. Assay buffer consisted of 25 mM MOPS (pH 7.4), 25 mM KCl, 1 mM EDTA, 0.01% fatty acid-free BSA, 0.01% Tween-20, 1 mM TCEP).

#### Binding Studies

An exemplar protocol for NCOR1/MED1 K_d_ tests was as follows: various His-PPARG-LBD (f.c. 2 nM) were diluted to 8x final concentration (f.c.) assay buffer. RXRA^WT^ or RXRA^S427F^ were diluted to 4x final concentration in assay buffer (f.c. 2 nM). Anti-6HIS Tb cryptate gold was diluted to 8x final concentration (f.c. 0.9 nM) in Terbium Detection Buffer. His-PPARG LBD dilutions and Anti-6HIS Tb cryptate gold were mixed 1:1 and incubated for 30 min at RT. Separately, NCOR1 or MED1 peptides were serially diluted to 2x final concentration in assay buffer with streptavidin-d2 with streptavidin-d2 held at a constant 2-fold excess to biotin at each respective concentration. Five microliters of PPARG/Ant-6HIS Tb cryptate gold, 5 µL RXRA, and 10 µL peptide mixtures were then mixed and incubated for at least 1 h at RT. HTRF signals were monitored using an Envision. Curves were fit using GraphPad software and a 4-parameter dose-response fit with Hill slopes constrained to 1.0.

#### Compound Studies

General protocols were as follows:

*NCOR1 recruitment*: Compound potency (EC_50_) and maximal extent of NCOR1 recruitment to PPARG were assessed using a TR-FRET binding assay measuring association of a biotinylated NCOR1 ID2 peptide (Biotin-GHSFADPASNLGLEDIIRKALMG-amide) to PPARG/RXRA LBD heterodimer. Specifically, 20 µL of TR-FRET master mix consisting of 2 nM WT PPARG LBD, 2 nM WT RXRA LBD or mutant S427F RXRA LBD, 50 nM NCOR1, 80 nM Rosiglitazone, 25 nM streptavidin-d2 (Cisbio) and 0.3 nM Anti-His Tb (Cisbio) in 25 mM MOPS pH 7.4, 25 mM KCl, 1 mM EDTA, 0.01% BSA, 0.01% Tween-20 and 1 mM TCEP was added to 384-well plates containing duplicate 10-point dose response titrations of compounds in 60 nL DMSO (0.3% f.c. DMSO (v/v)). Mixtures were incubated for 3 h and read in an EnVision plate reader (Perkin Elmer) with Ex/Em 615/665. To determine the potency (EC_50_) and extent of NCOR1 recruitment, TR-FRET ratios were normalized to the average ratio of DMSO control wells (0%) and to the average maximum ratio for positive control compound (T0070907; defined as 100%) in CDD Vault and analyzed using the Levenberg-Marquardt algorithm.

*MED1 blockade*: Compound potency (IC_50_) and maximal extent of MED1 repulsion to PPARG were assessed using a TR-FRET binding assay measuring association of a biotinylated MED1 LxxLL peptide (Biotin- VSSMAGNTKNHPMLMNLLKDNPAQ-amide) to PPARG/RXRA LBD heterodimer. Specifically, 20 µL of TR-FRET master mix consisting of 2 nM WT PPARG LBD (His-tagged), 2 nM WT or S427F RXRA LBD (FLAG-tagged), 350 nM NCOR1, 80 nM Rosiglitazone, 175 nM streptavidin-d2 (Cisbio) and 0.3 nM Anti-His Tb (Cisbio) in 25 mM MOPS pH 7.4, 25 mM KCl, 1 mM EDTA, 0.01% BSA, 0.01% Tween- 20 and 1 mM TCEP was added to 384-well plates containing duplicate 10-point dose response titrations of compounds in 60 nL DMSO (0.3% DMSO f.c. (v/v)). Mixtures were incubated for 3 h and read in an EnVision plate reader (Perkin Elmer) with Ex/Em 615/665. To determine the potency (IC_50_) and extent of MED1 repulsion, TR-FRET ratios were normalized to the average ratio of DMSO control wells (0%) and to the average minimum ratio for positive control compound (GW9662; defined as 100%) in CDD Vault and analyzed using the Levenberg-Marquardt algorithm.

#### Temperature-dependent rate studies

PPARG LBD and RXRA WT LBD (f.c. 2 nM), biotinylated NCOR1 (f.c. 50 nM), streptavidin-d2 (f.c. 300 nM), Anti-His Tb gold (f.c. 0.3 nM) were pre-equilibrated in assay buffer. Compound stocks (10 mM DMSO, 1% DMSO f.c.) were prepared in a 3-fold, 10 pt dilution scheme starting at top f.c. of 100 µM. Reactions were initiated by adding pre-equilibrated protein/peptide/FRET mixtures to compound for a final reaction volume of 20 µL in ProxiPlates. At various time points, FRET plates were read in an

Envision Plate reader. Time points collected were 5, 15, 45, 90, 180 and 360 minutes. 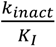 was determined by fitting the background subtracted TR-FRET ratio vs time to the one phase exponential association in Graphpad Software. Fitted k_obs_ values were then plotted against [Compound] (x) and fit to the following custom-entered equation in GraphPad software:

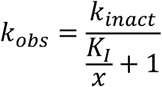

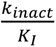 values were calculated from the resulting fit, plotted against temperature (K) and fit to the Eyring equation:

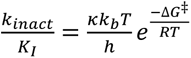; with:

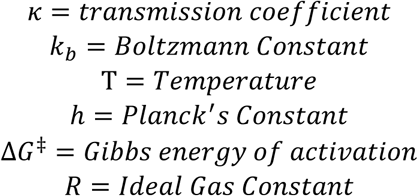

#### Covalent kinetic parameter determination

The general procedure for k_inact_ and K_I_ determination was as follows: 10x f.c. WT PPARG LBD in assay buffer and 10x f.c. Anti-6HIS-Tb gold in Terbium Detection Buffer were mixed 1:1 and incubated for 30 minutes at room temperature. The PPARG-antibody (f.c. 2 nM PPARG LBD and 0.9 nM Anti-6HIS Tb gold) mixture was mixed in assay buffer with Biotin-NCOR1 (f.c. 50 nM), streptavidin-d2 (f.c. 100 nM) and WT RXRA LBD (f.c. 2 nM). Compounds were diluted to 5x using a 3-fold, 10-point dilution scheme in assay buffer (1% DMSO f.c.). Eight microliters of protein/peptide/FRET mixtures were added to 2 µL of compound mixtures. Reactions were incubated at 25°C in the dark. At each time point, reactions were quenched by addition of Rosiglitazone (10 µM f.c.). Plates were then read at least 2 minutes after Rosiglitazone addition. Time points collected were 2, 5, 10, 15, 30, 60, 120 and 180 minutes. TR-FRET ratios were background subtracted (TR-FRET ratio – DMSO Control Signal = Background). Background subtracted TR-FRET ratio was then plotted over time at each inhibitor concentration and fit to the one phase exponential association in Graphpad Software. Fitted k_obs_ values were then plotted against [Compound] (x) and fit to the following custom-entered equation in GraphPad:

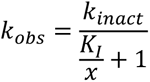

#### PPARA selectivity

NCOR1 recruitment: Compound potency (EC_50_) and maximal extent of NCOR1 recruitment to PPARA were assessed with a TR-FRET binding assay measuring association of a biotinylated NCOR1 ID2 peptide (Biotin-GHSFADPASNLGLEDIIRKALMG-amide) to PPARA/RXRA LBD heterodimer. Specifically, 10 µL of TR-FRET master mix consisting of 2 nM WT PPARA LBD, 2 nM WT RXRA LBD, 500 nM NCOR1, 1000 nM streptavidin-d2 (Cisbio) and 0.9 nM Anti-His Tb gold (Cisbio) in 25 mM MOPS pH 7.4, 25 mM KCl, 1 mM EDTA, 0.01% BSA, 0.01% Tween-20 and 1 mM TCEP was added to 384-well plates containing duplicate 10-point dose response titrations of compounds (1% f.c. DMSO (v/v)). Mixtures were incubated for 3 h and read in an EnVision plate reader (Perkin Elmer) with Ex/Em 615/665. To determine the potency (EC_50_) and extent of NCOR1 recruitment, TR-FRET ratios as a function of compound concentration were normalized by taking the ratio of the TR-FRET ratio observed at the respective concentrations relative to base line signal. The resulting fold change in TR-FRET ratio at each concentration was fitted and analyzed in GraphPad.

#### PPARD selectivity

NCOR1 recruitment: Compound potency (EC_50_) and maximal extent of NCOR1 recruitment to PPARD were assessed with a TR-FRET binding assay measuring association of a biotinylated NCOR1 ID2 peptide (Biotin-GHSFADPASNLGLEDIIRKALMG-amide) to PPARD/RXRA LBD heterodimer. Specifically, 10 µL of TR-FRET master mix consisting of 2 nM WT PPARD LBD, 2 nM WT RXRA LBD, 500 nM NCOR1, 1000 nM streptavidin-d2 (Cisbio) and 0.9 nM Anti-His Tb gold (Cisbio) in 25 mM MOPS pH 7.4, 25 mM KCl, 1 mM EDTA, 0.01% BSA, 0.01% Tween-20 and 1 mM TCEP was added to 384-well plates containing duplicate 10-point dose response titrations of compounds (1% f.c. DMSO (v/v)). Mixtures were incubated for 3 h and read in an EnVision plate reader (Perkin Elmer) with Ex/Em 615/665. To determine the potency (EC_50_) and extent of NCOR1 recruitment, TR-FRET ratios as a function of compound concentration were normalized by taking the ratio of the TR-FRET ratio observed at the respective concentrations relative to base line signal. The resulting fold change in TR-FRET ratio at each concentration was fitted and analyzed in GraphPad.

### Surface Plasmon Resonance Experiments (SPR)

SPR analysis was performed using an Cytiva S200 instrument at 25°C. N-terminal Avi-tagged PPARG LBD constructs were immobilized to an SA chip up to ∼1000RU using a running buffer consisting of 25mM MOPS pH 7.4, 25mM KCl, 1mM EDTA, 0.01% Tween 20, 1mM TCEP, and 1% DMSO. PPARG was incubated with FTX-6746 (or DMSO) for 1hr at room temperature to ensure complete labeling prior to immobilization. MED1 peptide serial dilutions (top concentration of 25 µM; 2-fold dilution) were injected over the PPARG surfaces at 30 µL/min for 60 sec with a 60 sec dissociation time. Report points at the end of the injection were plotted against MED1 concentration and fit to a Steady State Affinity model using Biacore Insights software. Percent activity was calculated by dividing the fitted R_max_ for each dose response by the theoretical R_max_ derived from ligand and analyte molecular weights.

### Hydrogen-Deuterium Exchange (HDX)

PPARG LBD and DMSO, T0070907 or FTX-6746 were incubated at 21°C for 3h. Samples were exchanged in D_2_O-containing HDX buffer and incubated at 4°C for 0, 10, 30, 60, 900 and 3600 seconds. Exchange was quenched with urea in acidic solution and samples were digested in pepsin solution at pH 2.5, 0°C, for 3 minutes. Digested peptides were desalted online at 1°C for approximately 2 minutes and subjected to HPLC-MS/MS analysis at 1°C. Differentials were then determined and analyzed in HDX WorkBench. See Supplementary Table 3 for technical and buffer specifications.

### Chemical synthesis – see supplemental methods

#### Cell Lines

The UC cell line UMUC9 was purchased from Millipore Sigma (08090505-1VL) and licensed by XimBio for research use. The UC cell lines HT1197 (CRL1473), 5637 (HTB-9), HT1376 (CRL-1472), J82 (HTB- 1), RT4 (HTB-2) and SW780 (CRL-2169) were purchased from ATCC. The UC cells lines KMBC2 (JCRB1148) and T24 (JCRB0711) were purchased from JCRB. The UC cell lines RT112, SCaBER, TCCSUP, UBLC1 and UMUC3 were supplied by WuXi AppTec for use.

#### Isogenic cell line generation

HT1197 cells were maintained in RPMI-1640 with 10% FBS (Gibco) at 37°C with 5% CO_2_. Cells were modified by CRISPR/Cas9 via electroporation of ribonucleoprotein particles generated with the following guide sequence: CTGCCAGTTTCGCTCCGTGG to generate a homozygous PPARG-C313A (TGC>GCC) substitution (Synthego). Cells were single-cell cloned and homozygous knock-in clone was verified by Sanger sequencing.

#### Pharmacodynamic cellular assays

For UMUC9 cells, analysis of relative levels of *IL1B* mRNA transcripts after 24h of treatment with DMSO or increasing concentrations of FX-909 was performed using qRT-PCR. A seven-point dose titration curve of FX-909 was utilized with starting and final concentrations of 100 nM and 0.1 nM, respectively. For HT1197 cells, analysis of relative levels of *ANGPTL4* mRNA transcript levels after treatment with DMSO or increasing concentrations of FX-909 was performed using quantitative RT-PCR. A seven-point dose titration curve of FX-909 was utilized with starting and final concentrations of 30 nM and 0.03 nM, respectively. Cells were processed for qRT-PCR analysis using Cells-to-C_T_ according to the manufacturer’s instructions for 2-step workflow of reverse transcription and qPCR following cell lysis (Thermo Fisher). For HT1197 cells, *ANGPTL4* and *TBP* (housekeeping) mRNA levels were quantitated and for UMUC9 cells, *IL1B* and *TBP* (housekeeping) mRNA levels were quantitated. Probe-based quantitative PCR on the ABI QuantStudio 7 Flex machine was performed, and data was analyzed using comparative C_T_ method (ΔΔC_T_). IC_50_ values were calculated using GraphPad Prism 9.

#### Cellular viability assays

UC cell lines were plated in 6-well plates at the densities indicated in Table 1 and incubated overnight. The following day, FX-909, FTX-6746 or T0070907 was added to the cells with triplicate wells for each dose. Cells were incubated at 37°C in 5% CO_2_ for 11 to 14 days as indicated in Table 1. For cell lines with an assay length of 11 days, medium and compound were replaced with fresh components on days 4 and 7. For cell lines with an assay length of 14 days, medium and compound were replaced with fresh components on days 4, 7 and 11. On the last day, cells were washed with PBS and stained with crystal violet solution (0.5% crystal violet in 20% methanol) for 10 minutes. Cells were then washed two times with PBS and allowed to dry. Imaging and quantitation of cell density were performed using the Licor Odyssey or absorbance measurement at 565 nm. Relative GI_50_ values were calculated using GraphPad Prism 9 by defining DMSO treated wells as 100% growth.

**Table 1.**
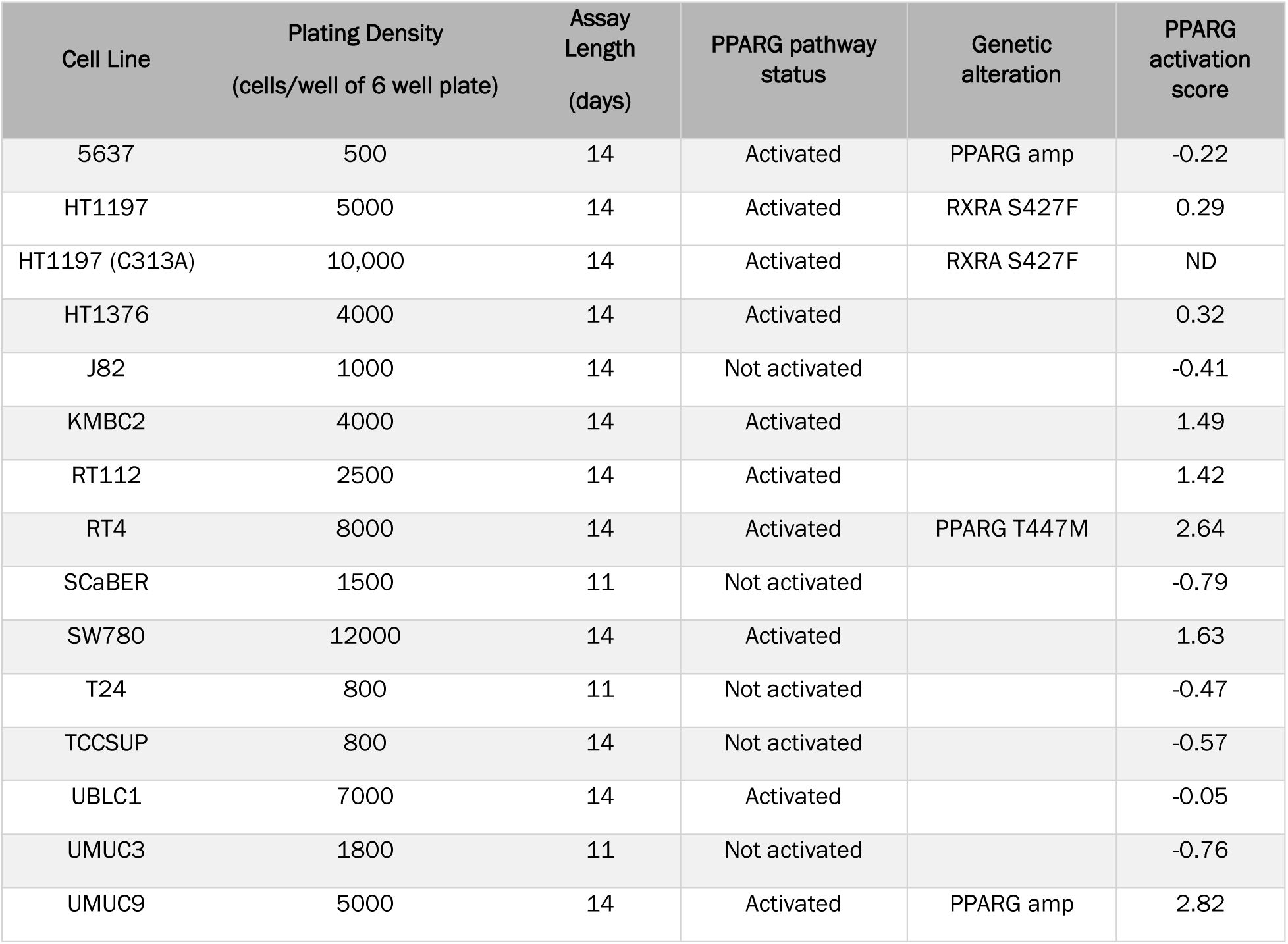

| Cell Line | Plating Density<br>(cells/well of 6 well plate) | Assay Length<br>(days) | PPARG pathway status | Genetic alteration | PPARG activation score |
| --- | --- | --- | --- | --- | --- |
| 5637 | 500 | 14 | Activated | PPARG amp | -0.22 |
| HT1197 | 5000 | 14 | Activated | RXRA S427F | 0.29 |
| HT1197 (C313A) | 10,000 | 14 | Activated | RXRA S427F | ND |
| HT1376 | 4000 | 14 | Activated |  | 0.32 |
| J82 | 1000 | 14 | Not activated |  | -0.41 |
| KMBC2 | 4000 | 14 | Activated |  | 1.49 |
| RT112 | 2500 | 14 | Activated |  | 1.42 |
| RT4 | 8000 | 14 | Activated | PPARG T447M | 2.64 |
| SCaBER | 1500 | 11 | Not activated |  | -0.79 |
| SW780 | 12000 | 14 | Activated |  | 1.63 |
| T24 | 800 | 11 | Not activated |  | -0.47 |
| TCCSUP | 800 | 14 | Not activated |  | -0.57 |
| UBLC1 | 7000 | 14 | Activated |  | -0.05 |
| UMUC3 | 1800 | 11 | Not activated |  | -0.76 |
| UMUC9 | 5000 | 14 | Activated | PPARG amp | 2.82 |

#### Mass spectrometry (MS) assays

UC cell lines were plated in 10 cm tissue culture dishes (3 x 10^6^ cells/dish) 24 h prior to treatment. FX-909 treatment for 30 min was performed and cells were incubated at 37°C in tissue culture incubator. Following treatment, cells were harvested and pelleted by centrifugation (300 x *g*, 4 min at 4°C), washed twice with 1X PBS and transferred to 1 mL low retention Eppendorf tubes. Frozen cell pellets were lysed using 1X PBS (pH 7.4) + 0.5% NP-40. Samples were further homogenized, and DNA was sheared using sonication with a probe sonicator. Total protein was determined using a BCA assay and cell lysates were used immediately for each experiment. Total cell extract (50 μg) was aliquoted for each Tandem Mass Tag (TMT) channel for further downstream processing. Cell extracts were treated with 500 μM of Desthiobiotin (DBIA) for 1 h in the dark at room temperature. Excess DBIA, along with disulfide bonds were quenched and reduced using 5 mM dithiothreitol for 30 min in the dark at room temperature. Subsequently, reduced cysteine residues were alkylated using 20 mM iodoacetamide for 30 min in the dark at room temperature. To facilitate removal of quenched DBIA and incompatible reagents, proteins were precipitated using chloroform/methanol. Protein pellets were re-solubilized in 200 mM 4-(2-hydroxyethyl)-1-piperazinepropanesulfonic acid (EPPS) at pH 8.5 and digested using LysC and trypsin (1:100, enzyme to protein ratios) overnight at 37°C. The next day samples were labeled with TMT reagents and cysteine-containing peptides were enriched using Pierce streptavidin magnetic beads. MS data were acquired using an Orbitrap Fusion Eclipse mass spectrometer in-line with a Vanquish Neo LC system. Peptides were separated using an in-house 100 µm capillary column packed with 35 cm of Accucore 150 resin (2.6 μm, 150 Å) using 210 min gradients from 4 to 24% acetonitrile in 0.125% formic acid per run. Eluted peptides were quantified using high-resolution MS2 acquisition.

All acquired data were searched using the open-source Comet algorithm (release_2019010) using a previously described informatics pipeline^30–33^. Spectral searches were done using a custom FASTA-formatted database which included common contaminants, reversed sequences (Uniprot Human, 2021) and the following parameters: 50 PPM precursor tolerance, fully tryptic peptides, fragment ion tolerance of 0.9 Da and a static modification by TMTPro (+304.2071 Da) on lysine and peptide N termini. Carbamidomethylation of cysteine residues (+57.021 Da) was set as a static modification while oxidation of methionine residues (+15.995 Da) and DBIA on cysteine residues (+239.262) was set as a variable modification. Peptide spectral matches were filtered to a peptide and protein false discovery rate (FDR) of less than 1% using linear discriminant analysis employing a target-decoy strategy. For quantification of each MS2 spectrum, a total sum signal-to-noise of all reporter ions of 200 was required. Lastly, peptide quantitative values were normalized so that the sum of the signal for all proteins in each channel was equal to account for sample loading differences (column normalization). TMT reporter ion sum-signal-to-noise for each experiment was used to calculate the competition ratios by dividing the control channel (DMSO) by the electrophile treated channel. Replicate measurements were averaged and reported as a single entry. Finally, cysteine site engagement was normalized to matched full proteomes to ensure changes were not due to differential protein expression.

#### RNA sequencing and analysis

*RNA sequencing.* UMUC9, HT1197 and 5637 cells were seeded in 6-well dishes at 5 x 10^5^ cells/well, 5 x 10^5^ cells/well and 2.5 x 10^5^ cells/well respectively. After 24 h, cells were treated in quadruplicate with DMSO, 0.5 μM FX-909, FTX-6746 or T0070907. After treatment, cells were washed 2X with PBS and lysed on the plate with 350 μl Buffer RLT from Qiagen RNeasy Plus Mini kit (Qiagen Cat No. 74134). RNA was extracted following Qiagen RNeasy Mini Kit instructions. RNA was quantified using Nanodrop One Spectrophotometer (Thermo Scientific). 1 μg of RNA at 50 ng/μl was used to construct total mRNA libraries using poly A enrichment and sequencing using Illumina Novaseq PE150bp reads (Novogene Corporation Inc, Sacramento, CA). Raw reads were mapped to GRCh38 with HISAT2 v2.0.5 using default parameters and gene-level quantification was performed with featureCount v1.0.5. All downstream data analyses were performed in R programming language.

*QC and outlier detection.* Low-expressing genes were filtered out based on read counts using the *edgeR* ‘filterByExpr’ function. The counts per million (CPM) matrix was normalized using the *edgeR* TMM method. Principal component analysis (PCA) was performed on normalized expression data for each cell line. One sample (5637 cell line treated with DMSO, replicate 3) out of the total of 48 samples was identified as an outlier and removed from analysis.

*PCA of gene expression changes.* After outlier removal, per-sample, per-gene log2(fold change) values were calculated by subtracting the mean of DMSO samples per cell line in the log2-transformed TMM-normalized CPM matrix. After removing genes whose expression changes have a variance of less than 0.05, PCA was performed on the log2(fold change) matrix.

*Differential expression*. After outlier removal, low-expressing genes were filtered out based on read counts using the *edgeR* ‘filterByExpr’ function. Data were normalized using the *DESeq2* VST method. Genes whose expression changes have a variance of less than 0.05 (see above) were further filtered out. The *lmerSeq* package ^34^ was used to fit a linear mixed effects model with treatment (FX-909, FTX- 6746, T0070907, or DMSO) specified as a fixed effect and cell line (UMUC9, HT1197, or 5637) as a random effect, as described by the following model design:

*Gene_i_ ∼ Treatment + (1|Cell_Line)*

Treatment-induced gene expression changes were derived by evaluating the following contrasts in the mixed effect model: FX-909 vs DMSO, FTX-6746 vs DMSO, and T0070907 vs DMSO (Supplementary Data 1). Differentially expressed genes (DEGs) were defined as having FDR<0.05 and absolute log2 fold change greater than 0.6. Hierarchical clustering of DEGs was performed using the *ComplexHeatmap* package with ‘ward.D2’ clustering method and ‘euclidean’ distance.

*Pathway analysis*. GSEA was performed on differential expression analysis results from the FX-909 vs DMSO comparison. Genes were ranked based on *log2(FC)*(-log10(FDR))*, and GSEA was performed on the pre-ranked gene list using the *fgsea* ^35^ (v1.24.0) and *msigdbr* (v7.5.1) packages. GSEA results are reported in Supplementary Data 2.

#### Cut&Run analysis

Cut&Run was performed using the Epicypher CUTANA ChIC/CUT&RUN Kit V2.1 following the manufacturer’s recommendations. UMUC9 cells were plated in 15 cm plates, after 24 h cells were treated with DMSO or 0.5 µM FTX-6746 for 24 h. Cells were collected via trypsinization and counted with the Countess II Cell Counter (Life Technologies). 500,000 cells were used per Cut&Run reaction.

0.5 μg of each antibody was used: PPARG antibody (ThermoFisher A3409A-00 Lot A-2), RXRA Cell Signaling 3085 Lot 4, IgG Control from Epicypher kit. Cut DNA was prepared following the kit’s protocol. Libraries were prepared using the NEBNext® UltraTM II DNA Library Prep Kit for Illumina (#E7103S/L) with NEBNext Multiplex Oligos for Illumina (NEB) as per manufacturer’s instructions to select for short DNA fragments as outlined by the Epicypher CUTANA handbook. 2 ng of Cut&Run DNA was used to prepare each library. Libraries were quantified using Qubit dsDNA HS reagents (ThermoFisher Scientific) and the TapeStation D1000 (Agilent) and equimolar amounts of each sample were pooled and sequenced via paired-end 50 base pair reads on an Illumina NextSeq (Novogene). Samples were de-multiplexed (Novogene) and Fastq files were uploaded to BasePair for analysis using CUT&RUN QC, Alignment (Bowtie2) and CUT&RUN Peaks, Motif (MACS2, Homer) pipelines with default MACS2 peak calling parameters. Differential peaks were called with DiffBind (Bioconductor) and heatmap created with DeepTools with a window of 5000 bp upstream and downstream of annotated TSS.

PPARG activation gene signature derivation. Starting with a previously reported list of 197 genes significantly deregulated by PPARG siRNA knockdown^36^, DESeq2^37^ was used to determine differentially expressed genes between the 25% highest PPARG expressing and the 25% lowest PPARG expressing primary bladder cancer tissues in the TCGA (n=407). The resulting overlapping 120 differentially expressed genes (Supplementary Figure 13A) were evaluated for biologically concordant regulation (Supplementary Figure 13B), and a final list of 100 genes excluding PPARG (54 upregulated by PPARG and 46 downregulated by PPARG) were included in the proposed PPARG activation gene signature.

PPARG activation score calculation. CCLE mRNA expression data [n=1,517 cell lines; protein-coding genes only; log2(TPM+1)] were quantile normalized. Expression Z-score was calculated for each gene across all cell lines. 93 out of the 100 PPARG activation genes were present in the CCLE mRNA expression data (51 upregulated; 42 downregulated by PPARG). Coherence of the expression of these genes was assessed in the selected 14 UC cell lines: unbiased hierarchical clustering showed that 95% of the genes (88/93) clustered within their expected groups of PPARG up- or downregulated genes (Table 1; Supplementary Figure 13C). PPARG activation score for each cell line was calculated using the formula:

PPARG activation score = mean (Z-score of UP genes) – mean (Z-score of DOWN genes) PPARG activation scores are reported in Table 1.

#### Data availability

TCGA mRNA expression data were downloaded from the UCSC Toil RNAseq Recompute Compendium^38^ (‘TcgaTargetGtex_gene_expected_count.gz’ file, v2016-09-03). CCLE mRNA expression data were downloaded from the DepMap 24Q2 Public release (DepMap, Broad, 2024; ‘OmicsExpressionProteinCodingGenesTPMLogp1BatchCorrected.csv’ file)^39^ .

### In vivo studies

#### Xenograft Efficacy Studies

##### UMUC9 xenograft study

NOD CRISPR Prkdc IL2r gamma (NCG) 6–8-week-old female mice were inoculated subcutaneously in the right flank with 5 x 10^6^ UMUC9 cells in 50% Matrigel in base medium. Animals were randomized in groups of 10 based on tumor size once tumors reached approximately 135 mm^3^. Animals were dosed with vehicle, FTX-6746 or FX-909 by oral gavage (PO) twice a day (BID) for 21 days. Animals were routinely assessed for any clinical changes, tumor size by caliper measurement, and body weight twice weekly. Following drug treatment, a subset of animals was observed for an additional 45 days with tumor and body weight measurements twice weekly.

##### Adipose tissue analysis

Inguinal white adipose tissue was collected from a subset of mice at day 21 from the UMUC9 xenograft efficacy study. The tissues were fixed in formalin and embedded in paraffin. FFPE blocks were sectioned and stained with H&E to allow visualization of the architecture of the adipose tissue.

##### HT1197 xenograft study

BALB/c nude 6-8-week-old female mice were inoculated subcutaneously in the right flank with 5 x 10^6^ HT1197 cells in 50% Matrigel in PBS. Animals were randomized in groups of 8 based on tumor size once tumors reach approximately 170 mm^3^. Animals were dosed with vehicle,

FTX-6746 or FX-909 by oral gavage for 42 days. Animals were routinely assessed for any clinical changes as well as tumor size by caliper measurement and body weight twice weekly.

##### Xenograft PK/PD Study

NCG 6-8-week-old female mice were inoculated subcutaneously in the right flank with 5 x 10^6^ UMUC9 cells in 50% Matrigel in base medium. Animals were randomized into groups of 5 based on tumor size once tumors reached 400-500 mm^3^. Animals were dosed with vehicle, FTX- 6746 or FX-909 by oral gavage BID for 3 doses. Eight h after the 3^rd^ dose, animals were sacrificed for collection of tumor and plasma samples for analysis. RNA was extracted from tumors and used for qPCR analysis of *IL1B* which was normalized with housekeeping genes (*GAPDH* and *TBP*). *IL1B* expression relative to vehicle control samples was assessed. Plasma samples underwent bioanalytical assessment of compound levels to correlate with PD analysis.

## Acknowledgements

We thank our colleagues Michaela Bowden, Mitch Lazar, Stephen Frye, Steve McKnight and Fraydoon Rastinejad for their helpful conversations during the generation of this data. We also thank our colleagues at ChemPartner, WuXi AppTec and the Bioinformatics CRO for their contributions to the work presented here.

## Supplemental Figures

**Supplementary Figure 1.**
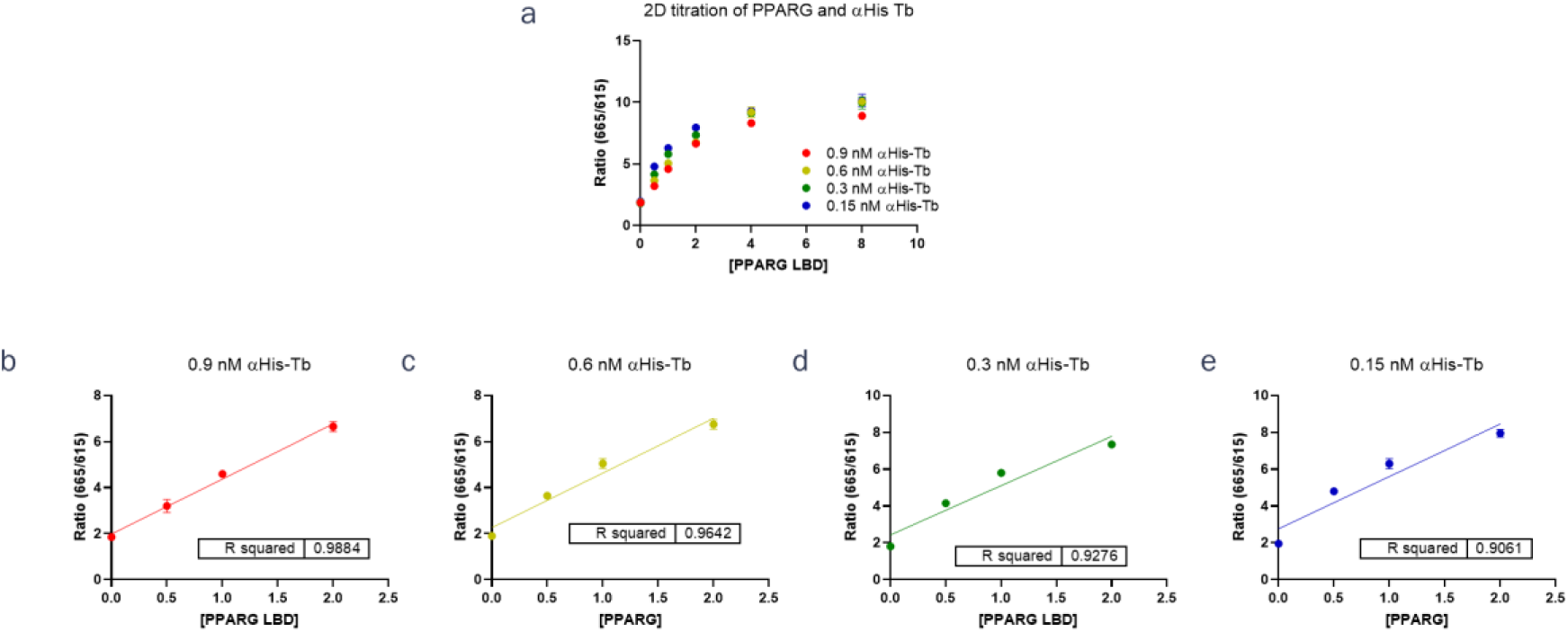
Assay optimization of PPARG TR-FRET assay for NCOR1 recruitment (a) 2D titration of PPARG and αHis-Tb antibody. (b-e) breakout of linear range with respect to [PPARG LBD] at individual αHis-Tb. Experiments were performed with 450 nM NCOR1 (∼K_d_) and streptavidin-d2. A concentration of 2 nM PPARG LBD was selected for subsequent experiments. Plots consist of the average of 2 technical replicates, error bars indicate the standard deviation of the mean.

**Supplementary Figure 2.**
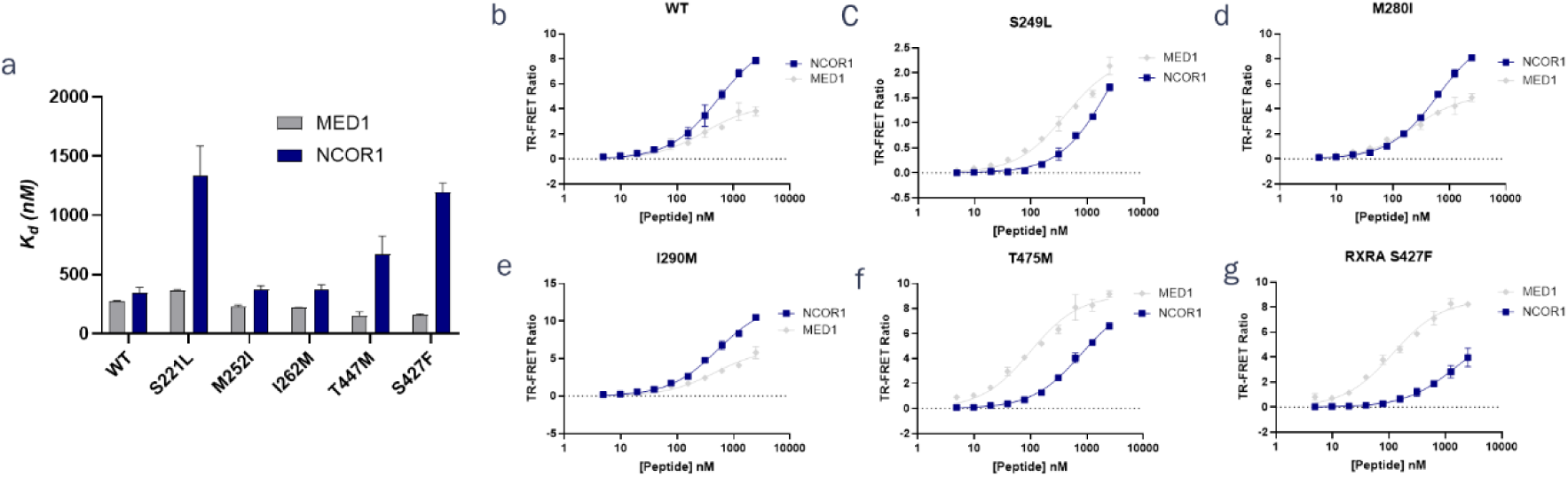
Biochemical affinity for MED1 and NCOR1 peptides of various MIUC mutants (a) K_d_ comparison of MIUC mutants for MED1 and NCOR1 K_d_ values represent the average of 2 biological replicates, each replicate performed in technical duplicate. Error bars indicate the S.E.M between the biological replicates (b-g) FRET binding curves for MIUC mutant binding to MED1 (gray) and NCOR1 (dark blue) peptides. Representative curves from 2 biological replicates are shown. Data points are the average of technical duplicates of a single biological replicate.

**Supplementary Figure 3.**
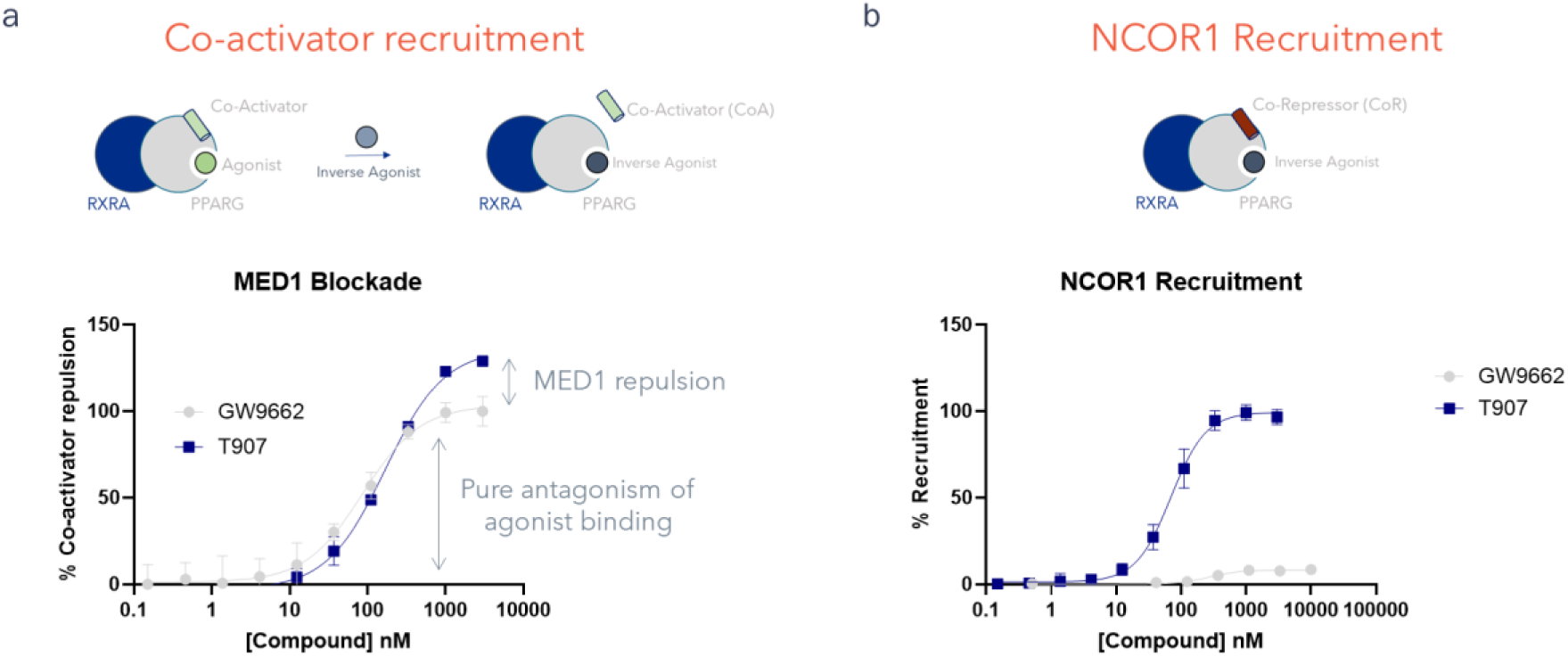
Assay overview for medicinal chemistry support. (a) Blockade of MED1 binding through agonist displacement without subsequent conformational biasing (GW9662, gray, neutral antagonism) and with subsequent conformational biasing (T0070907, Dark blue, inverse agonism). The 95% confidence interval of the EC_50_ values were 97 ± 38 nM for GW9662 and 178 ±39 nM for T0070907. Values are reported from a single biological replicate performed in technical duplicate (b) Recruitment of NCOR1 by T0070907 (dark blue, 95% C.I. of EC_50_ = 70 ± 2 nM) or GW9662 (gray, inactive). Values are reported from a single biological replicate performed in technical duplicate (a & b) The similar EC_50_ values for these compounds with divergent peptide recruitment profiles indicate a similar initial pose for binding and covalent reaction with subsequently divergent conformational biasing effects. Example curves are shown from a single biological replicate performed in technical duplicate. Curves are representatives of >10 biological replicates.

**Supplementary Figure 4.**
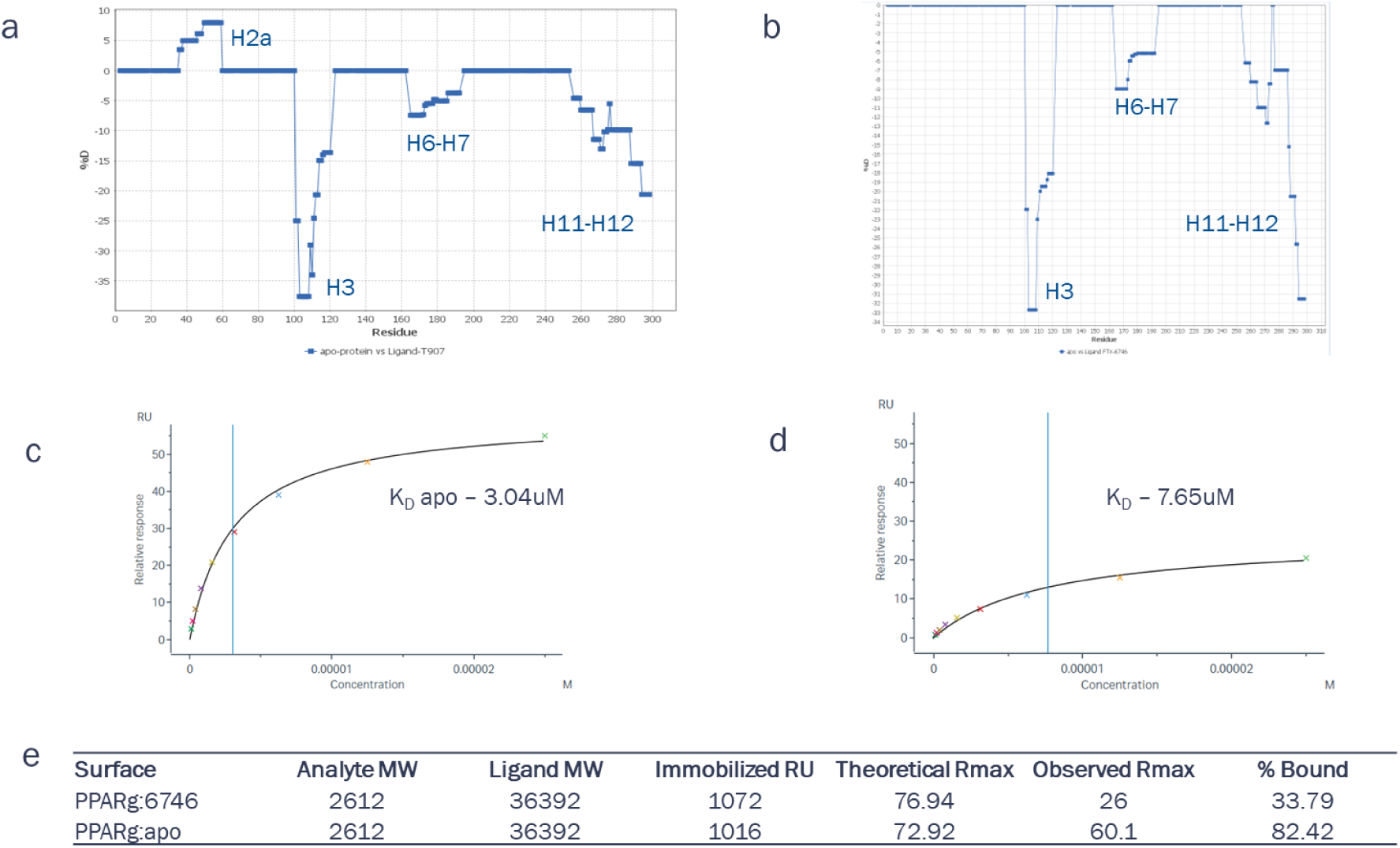
Biophysical validation of altered conformational sampling of PPARG. HDX shielding patterns of PPARG inverse agonists: (a) T0070907 relative to apo LBD. (b) FTX-6746 relative to apo LBD. The greater shielding in the H11-12 region seen with FTX-6746 indicates this compound more strongly stabilizes the inverse agonist conformation observed crystallographically. (c) and (d) SPR analysis of MED1 binding to (c) unmodified PPARG and (d) FTX-6746-modified PPARG. (e) Quantitative summary of SPR analyses. For both conditions, triplicate measurements were collected.

**Supplementary Figure 5.**
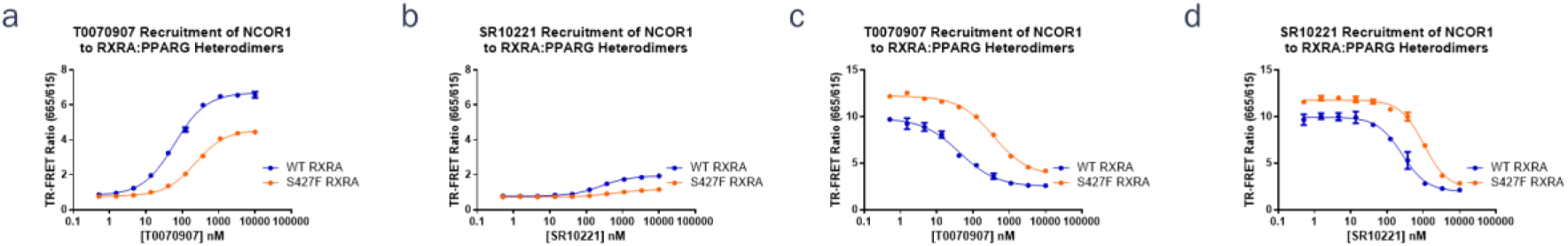
NCOR1 recruitment and MED1 blockade of literature tools compounds T0070907 and SR10221 with RXRA^WT^ or RXRA^S427^–containing PPARG heterodimers (a) T0070907 recruitment of NCOR1 to RXRA^WT^(blue) and RXRA^S427F^ heterodimers with PPARG (95% C.I. of EC_50_ values: RXRA^WT^ = 62 ± 8 nM and RXRA^S427F^ = 230 ± 20 nM). (b) SR10221 recruitment of NCOR1 to RXRA^WT^(blue) and RXRA^S427F^ heterodimers with PPARG (95% C.I. of EC_50_ values: RXRA^WT^ = 260 ± 40 nM and RXRA^S427F^ = 659 ± 130 nM). (c) T0070907 blockade of MED1 to RXRA^WT^(blue) and RXRA^S427F^ heterodimers with PPARG (95% C.I. of EC_50_ values: RXRA^WT^ = 45 ± 20 nM and RXRA^S427F^ = 370 ± 70 nM). (d) SR10221 blockade of MED1 to RXRA^WT^(blue) and RXRA^S427F^ heterodimers with PPARG (95% C.I. of EC_50_ values: RXRA^WT^ = 280 ± 80 nM and RXRA^S427F^ = 1,000 ± 230 nM). (a—d) Plots represent a single biological replicate, performed in technical duplicate for each measurement. Data are representative of ≥2 biological replicates.

**Supplementary Figure 6.**
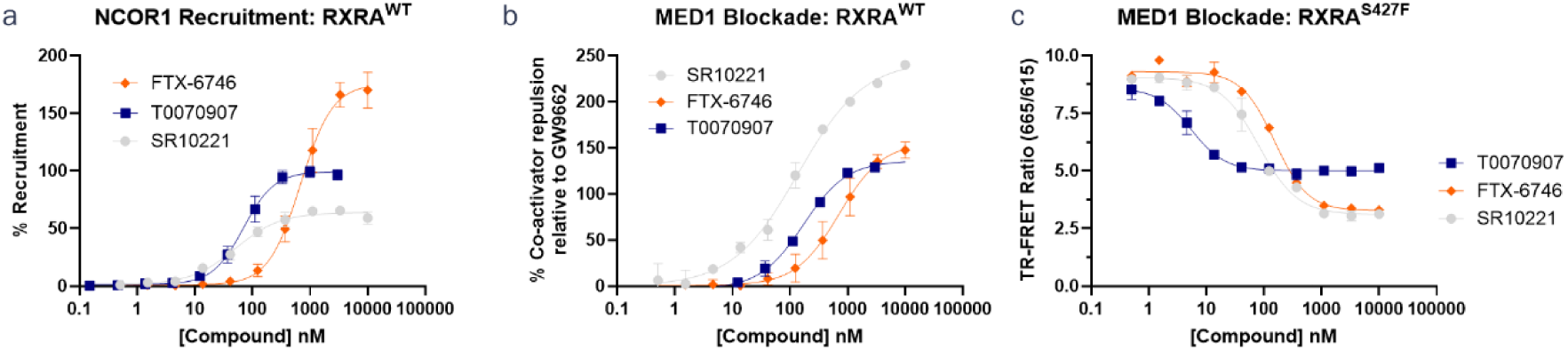
Comparison of FTX-6746, T0070907 and SR10221 in (a) NCOR1 recruitment with RXRA^WT^-containing heterodimers, 95% C.I. of EC_50_ values: SR01221 = 53 ± 16 nM; T0070907 = 70 ± 1.5 nM; FTX-6746 = 690 ± 70 nM; (b) MED1 blockade with RXRA^WT^-containing heterodimers and 95% C.I. of EC_50_ values: SR01221 = 150 ± 50 nM; T0070907 = 180 ± 40 nM; FTX-6746 = 740 ± 160 nM (c) MED1 blockade with RXRA^S427F^-containing heterodimers (no Rosiglitazone present). 95% C.I. of EC_50_ values: SR01221 = 90 ± 20 nM; T0070907 = 5.5 ± 1.7 nM; FTX-6746 = 160 ± 30 nM (a and b) Plots represent a single biological replicate, performed in technical duplicate for each measurement. Data are representative of ≥2 biological replicates.

**Supplementary Figure 7.**
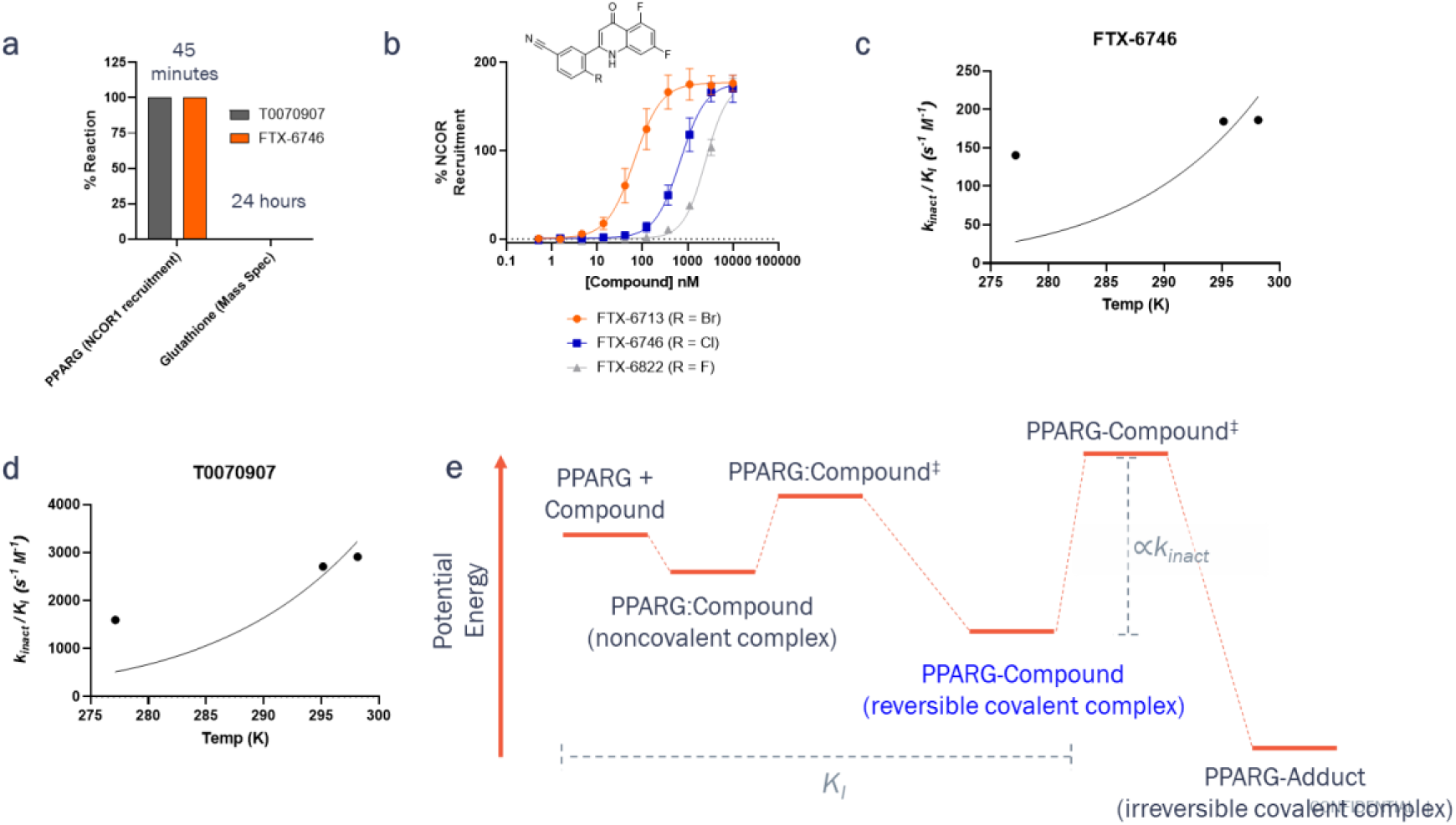
Mechanistic investigation of S_N_Ar capture of C313 (a) Relative reactivity of PPARG inverse agonists with PPARG C313 vs GSH.Percent reaction with PPARG is derived from the time point in which maximal FRET E_max_ was obtained using an NCOR1 recruitment assay. GSH reactivity was monitored by mass spec. NCOR1 recruitment assay were performed at room temperature while GSH reactivity was performed at 37°C. (b) Halogen SAR trend observed for PPARG inverse agonists. E_max_ remains unchanged among analogs as the final covalent adduct is the same while potency shows a reverse reactivity trend as is expected for S_N_Ar reactions. Data are from a single biological replicate measured in technical duplicate 95% C.I. of EC_50_ values: FTX-6713 = 70 ± 8 nM; FTX-6746 = 700 ± 70 nM; FTX-6822 = 5,000 ± 2,500 nM (c) Temperature-dependence of FTX-6746 rate constant determined through NCOR1-recruitment FRET assay. Trend is line the best fit of the data to the Eyring Equation. (d) Temperature-dependence of T0070907 rate constant determined through NCOR1- recruitment FRET assay. Trend is line the best fit of the data to the Eyring Equation. (e) Theoretical reaction coordinate rationalizing stepwise-nature of reaction mechanism and contextualizing relevant states measured by K_I_ and k_inact_ analyses. For (c) and (d), values from a single biological replicate performed in technical duplicate are shown. Data are representative of 3 biological replicates, each performed in technical duplicate.

**Supplementary Figure 8.**
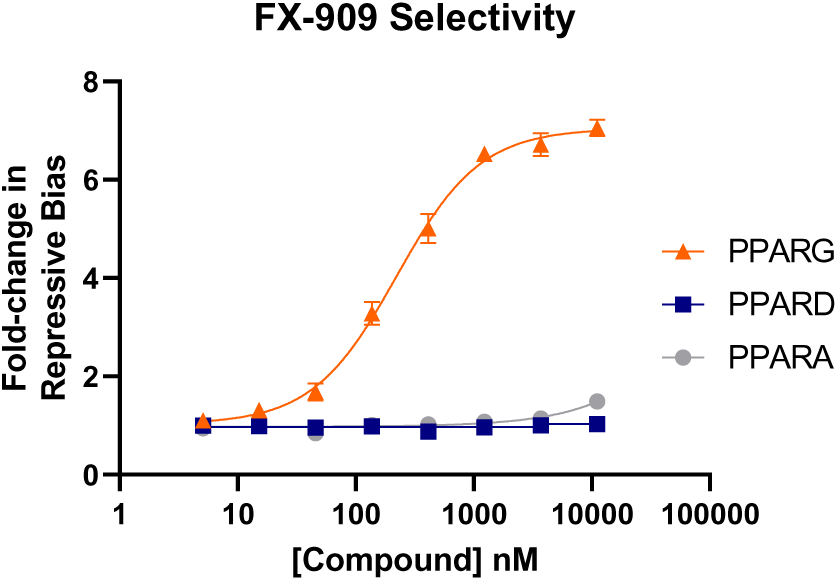
FX-909 selectivity as assessed by NCOR1 recruitment over PPARA and PPARD – Data are from a single biological replicate measured in technical duplicate 95% C.I. of EC_50_ values: PPARG = 200 ± 30 nM; PPARA > 100µM; PPARD > 100 µM.

**Supplementary Figure 9.**
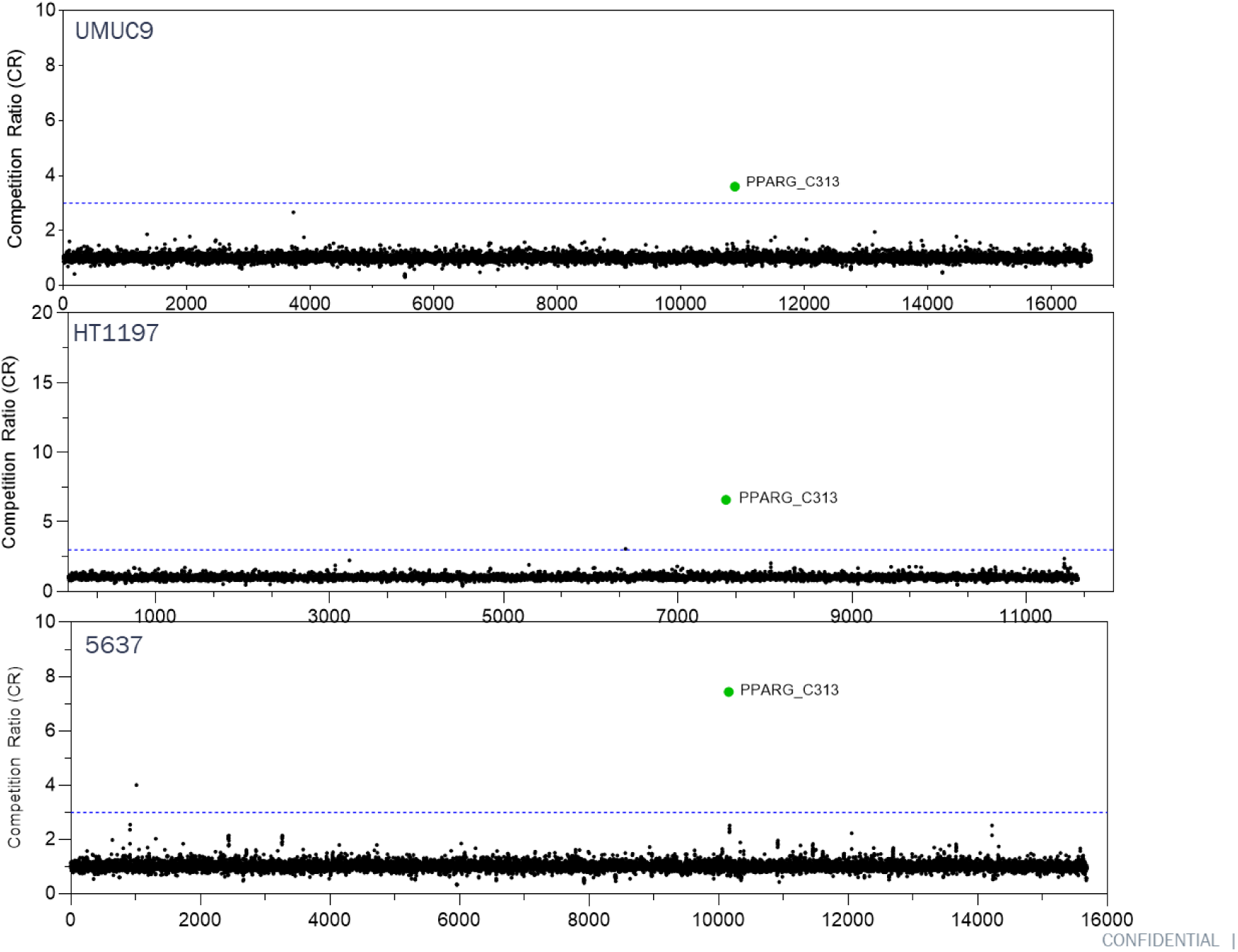
Selective engagement of FTX-6746 across urothelial carcinoma cell lines. Global cysteine profiling and engagement of FTX-6746 in UMUC9, HT1197 and 5637 cell lines following 30 min treatment. PPARG C313 was the only significantly engaged target (green). Competition ratio (CR) was calculated dividing the control channel (DMSO) by the electrophile treated channel. The blue dotted line indicates a CR of 3.

**Supplementary Figure 10.**
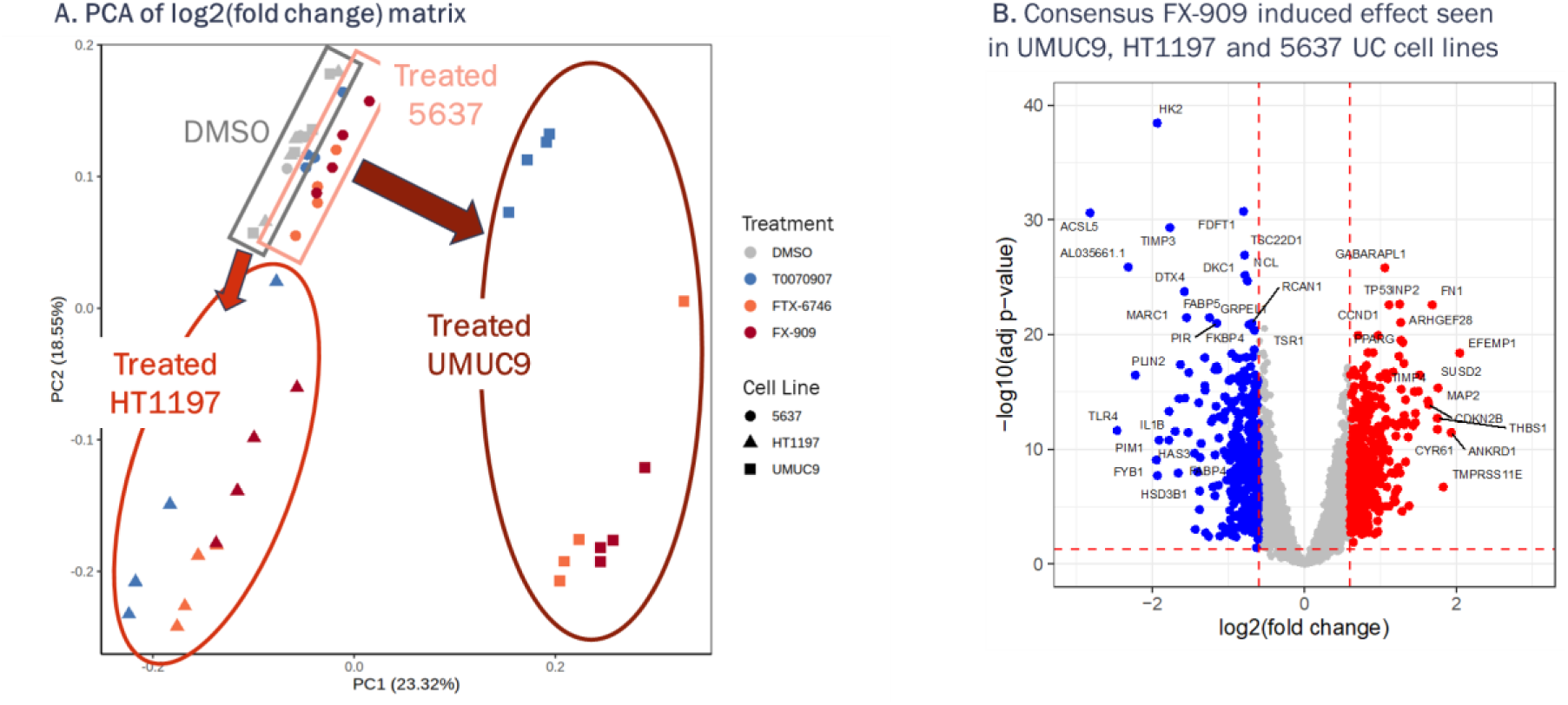
(A) Principal component analysis (PCA) of genome-wide expression changes (on log2 scale) induced by PPARG inverse agonists in UMUC9, HT1197 and 5637 cell lines after 24 hr treatment. (B) Consensus FX-909 induced differential gene expression was assessed using a mixed effect model where Treatment was a fixed effect and Cell Line was a random effect. Differentially expressed genes were determined by contrasting the FX-909 treatment with DMSO and using the cutoffs of FDR < 0.05 and absolute log2(fold change) > 0.6.

**Supplementary Figure 11.**
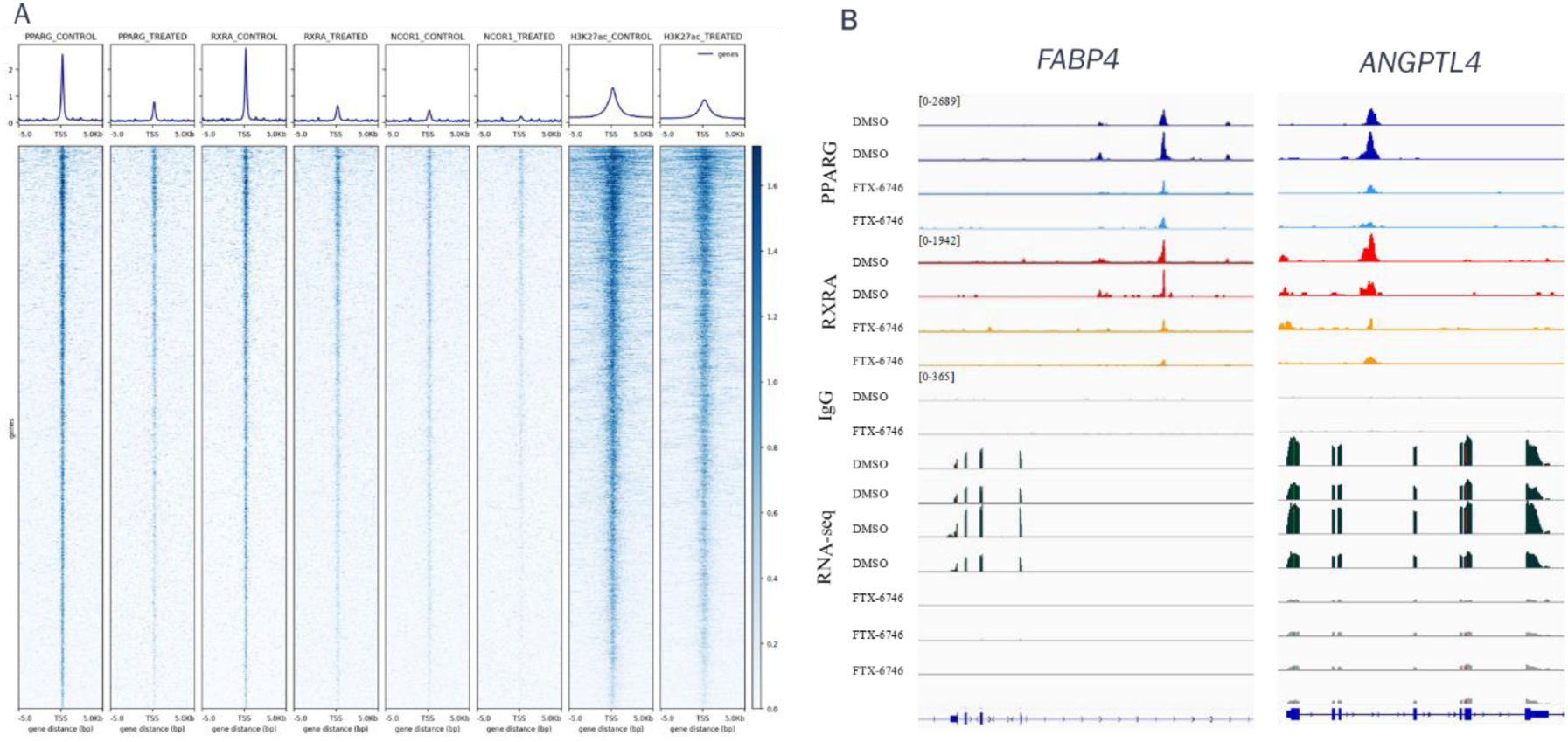
Genome-wide occupancy of PPARG, RXRA, NCOR1, and H327ac in UMUC9 treated with FTX-6746 by Cut&Run. UMUC9 cells were treated with DMSO or 0.5 µM FTX-6746 for 24 h followed by Cut&Run for the indicated proteins/histone modifications. (A) Heatmaps and metagene plots of PPARG, RXRA, NCOR1, and H3K27ac occupancy by Cut&Run in UMUC9 DMSO vs FTX-6746 treated samples. Heatmaps ranked by PPARG occupancy in DMSO treated control. (B) Example tracks of replicate Cut&Run and RNA-seq data for DMSO vs FTX-6746 treated UMUC9 cells at PPARG target gene loci, *FABP4* and *ANGPTL4*. Track snapshot created using IGV genome browser.

**Supplementary Figure 12.**
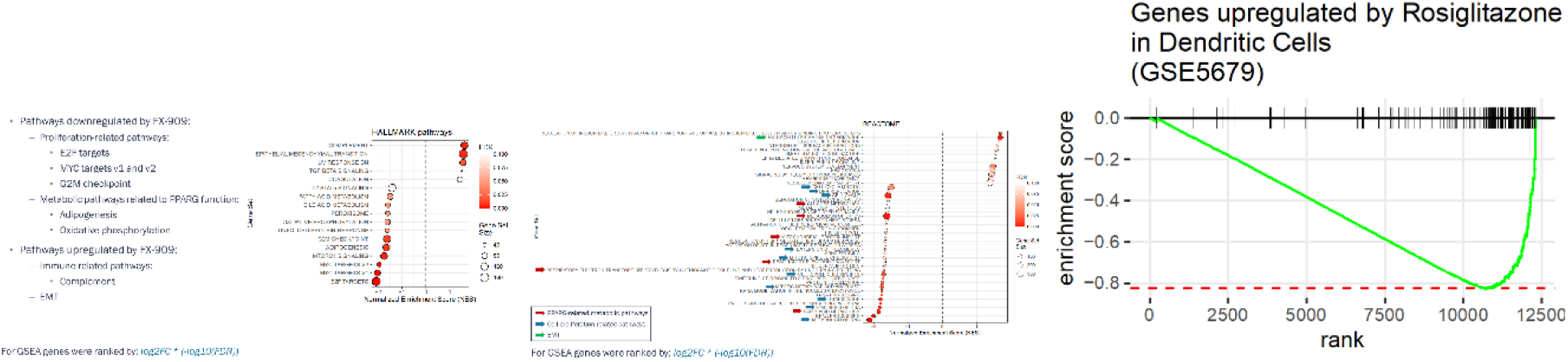
GSEA of the FX-909 induced gene expression changes. Genes were ranked by log2FC * (-log10(FDR)) and GSEA was performed using the Hallmark (A), Reactome, and ImmuneSigDB (C) gene set collections from the Broad’s MSigDB. (A and B) All pathways with FDR<0.1 are displayed. (C) Enrichment plot for a single gene set comprising of genes upregulated by the PPARG activator Rosiglitazone in dendritic cells.

**Supplementary Figure 13.**
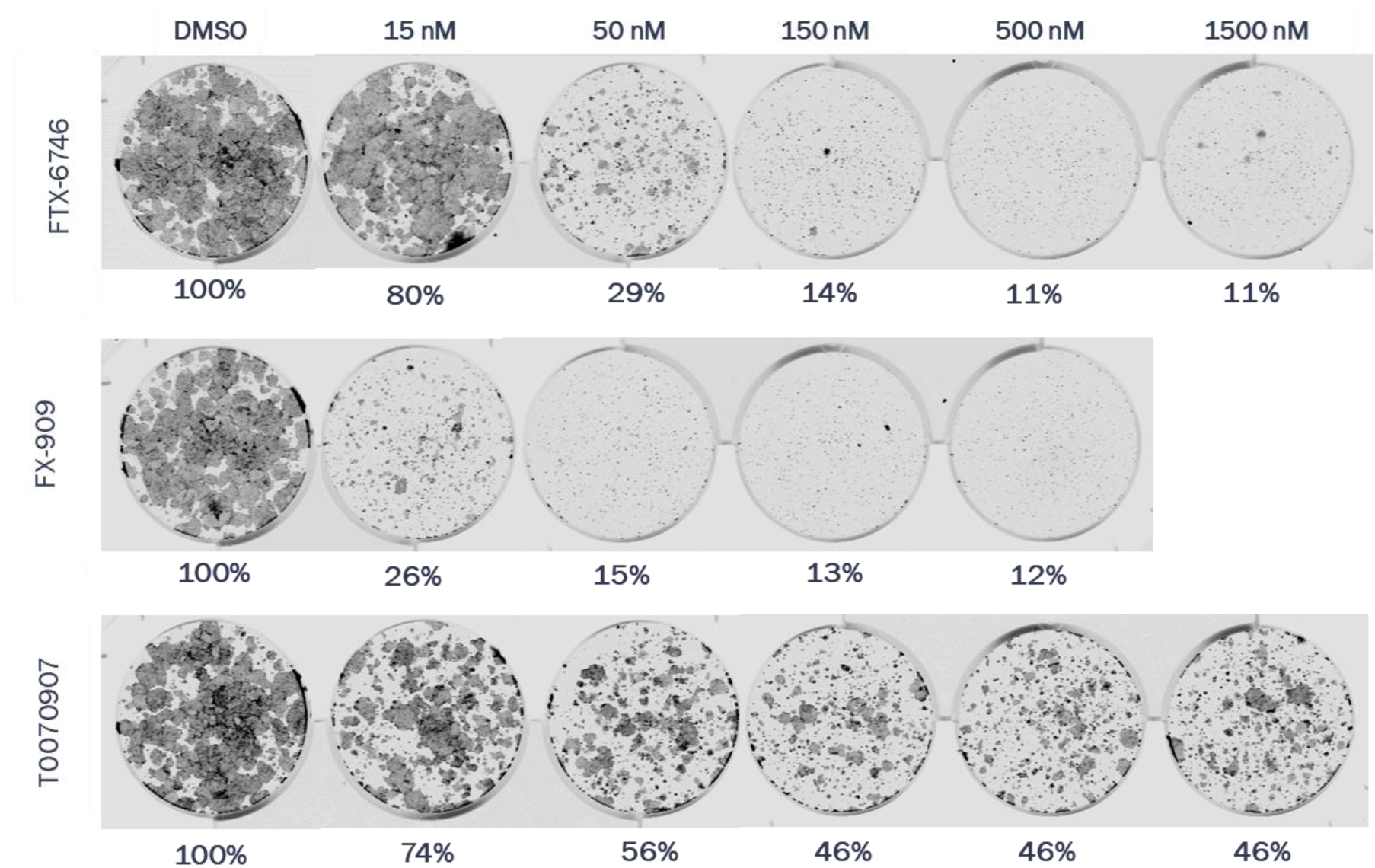
Clonogenic growth assay of HT1197 cells treated with FTX-6746, FX-909 or T0070907. Percent cell density relative to DMSO control well shown, data are representative of three independent experiments.

**Supplementary Figure 14.**
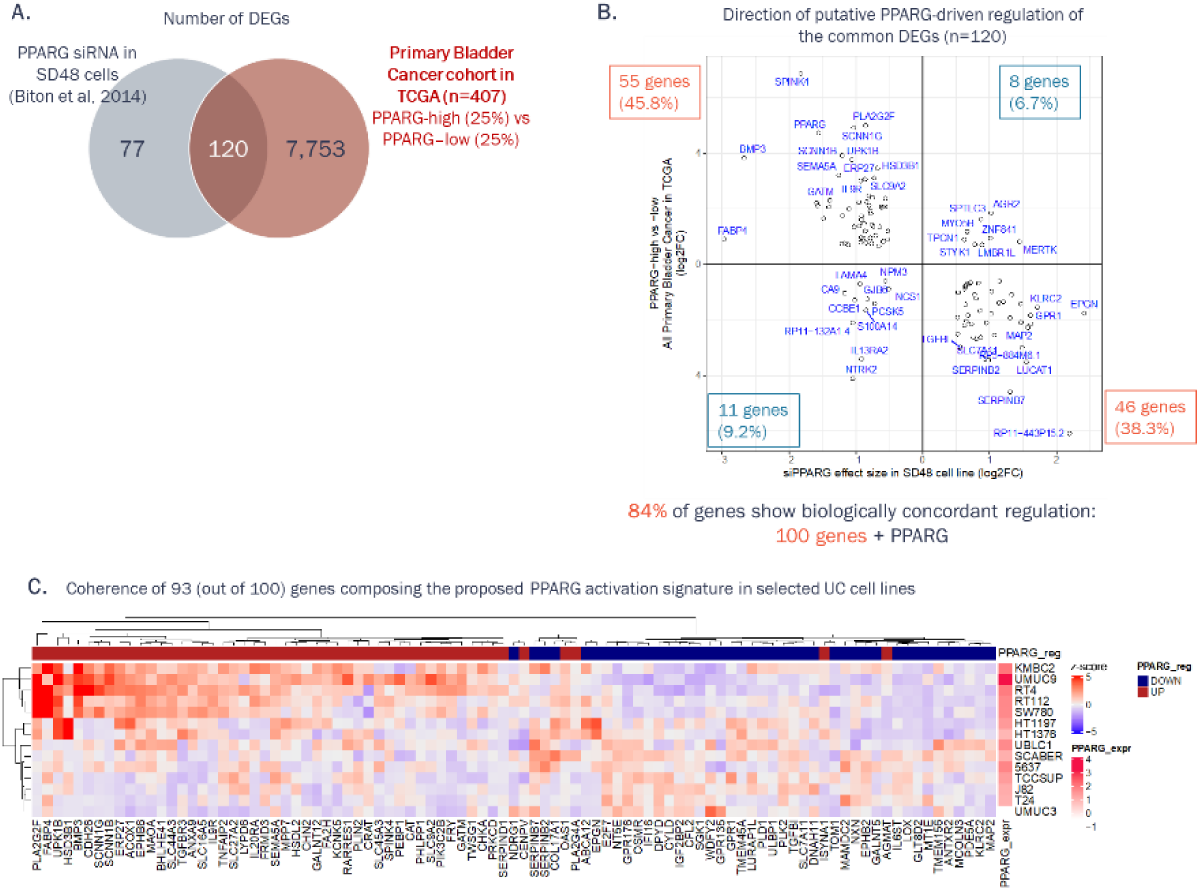
Derivation of PPARG activation gene signature. (A) Overlap between differentially expressed genes (DEGs) derived from PPARG siRNA knockdown^36^ and comparison of PPARG-high to PPARG-low primary bladder cancer tissues in the TCGA. (B) Evaluation of biologically concordant regulation of the common DEGs in the 2 data sets. Genes downregulated by PPARG knockdown are expected to be higher expressed in PPARG-high bladder cancer tissues (‘PPARG UP’ genes). Genes upregulated by PPARG knockdown are expected to be lower expressed in PPARG-high bladder cancer tissues (‘PPARG DOWN’ genes). (C) Hierarchical clustering of mRNA expression of the genes with biologically concordant regulation (i.e. final PPARG activation gene signature) in selected urothelial cancer (UC) cell lines from the CCLE.

**Supplementary Table 1.**
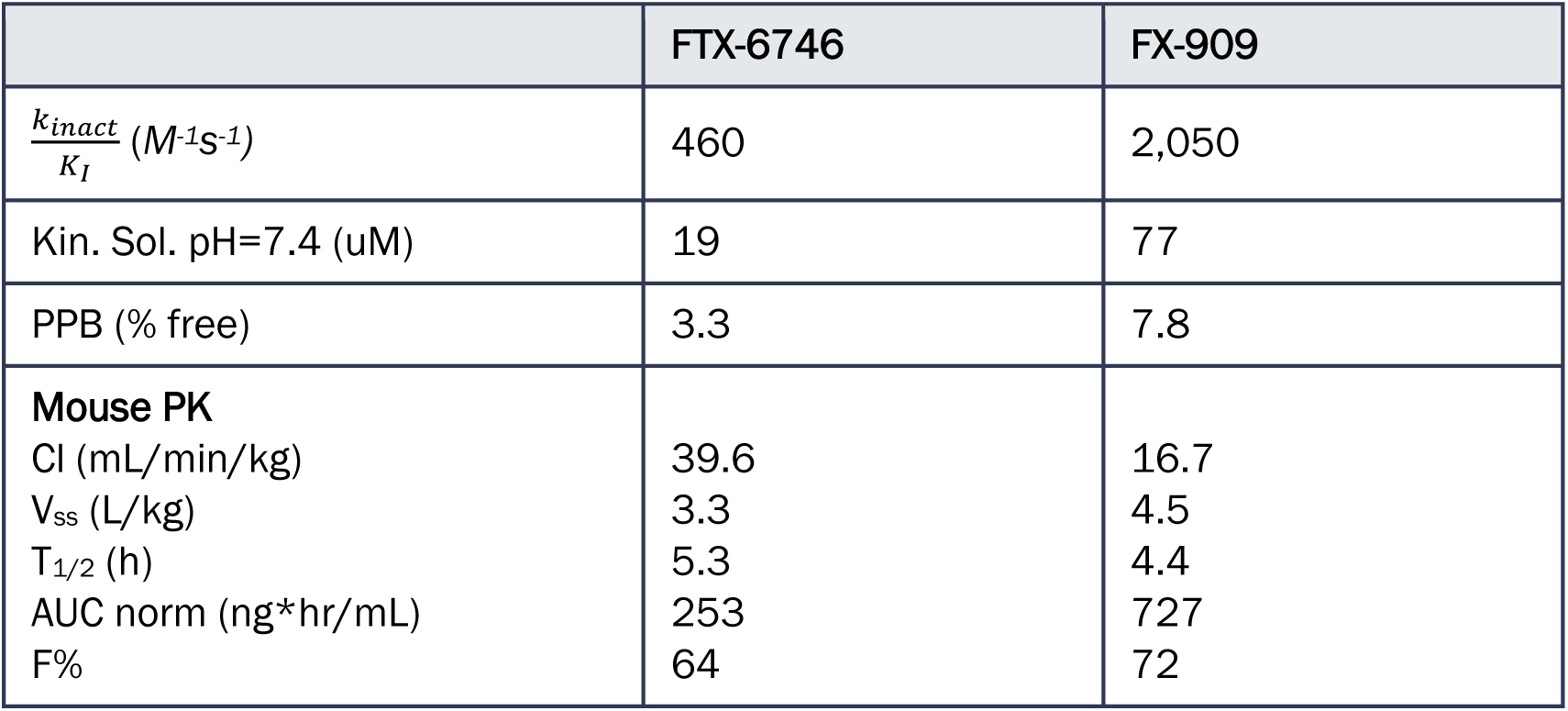
Comparison of FTX-6746 and FX-909.

**Supplementary Table 2.** Crystallographic Data Collection and Refinement Statistics.

| PPARG_cocrystal_FTX-6746 |  |
| --- | --- |
| **Data Collection** |  |
| Space group | *P1* |
| Cell dimensions |  |
| *a*, *b*, *c* (Å) | 53.13 70.33 92.68 |
| $\alpha$, $\beta$, $\gamma$ (°) | 101.19 106.58 104.98 |
| Resolution (Å) | 48.00 – 2.83 (2.98 – 2.83) |
| *R*<sub>merge</sub> | 0.074 (0.434) |
| *R*<sub>meas</sub> | 0.102 (0.620) |
| *R*<sub>pim</sub> | 0.074 (0.434) |
| *I* / $\sigma I$ | 9.1 (2.4) |
| *CC*1/2 | 0.995 (0.889) |
| Completeness (%) | 90.0 (83.7) |
| Redundancy | 3.7 (3.7) |
| **Refinement** |  |
| Resolution (Å) | 48.05 – 2.83 (2.98 – 2.83) |
| No. reflections | 22992 |
| *R*<sub>work</sub> / *R*<sub>free</sub> | 0.2462/0.3042 |
| No. atoms | 8190 |
| Protein | 8113 |
| Ligand | 63 |
| Water | 14 |
| *B*-factors | 90.09 |
| Protein | 96.24 |
| Ligand | 62.16 |
| Water | 55.02 |
| R.m.s. deviations |  |
| Bond lengths (Å) | 0.0066 |
| Bond angles (°) | 1.4042 |
| Ramachandran plot |  |
| Favored (%) | 98.3 |
| Allowed (%) | 1.2 |
| Outliers (%) | 0.5 |
| Statistics for the highest-resolution shell are shown in parentheses. |  |

**Supplementary Table 3.**
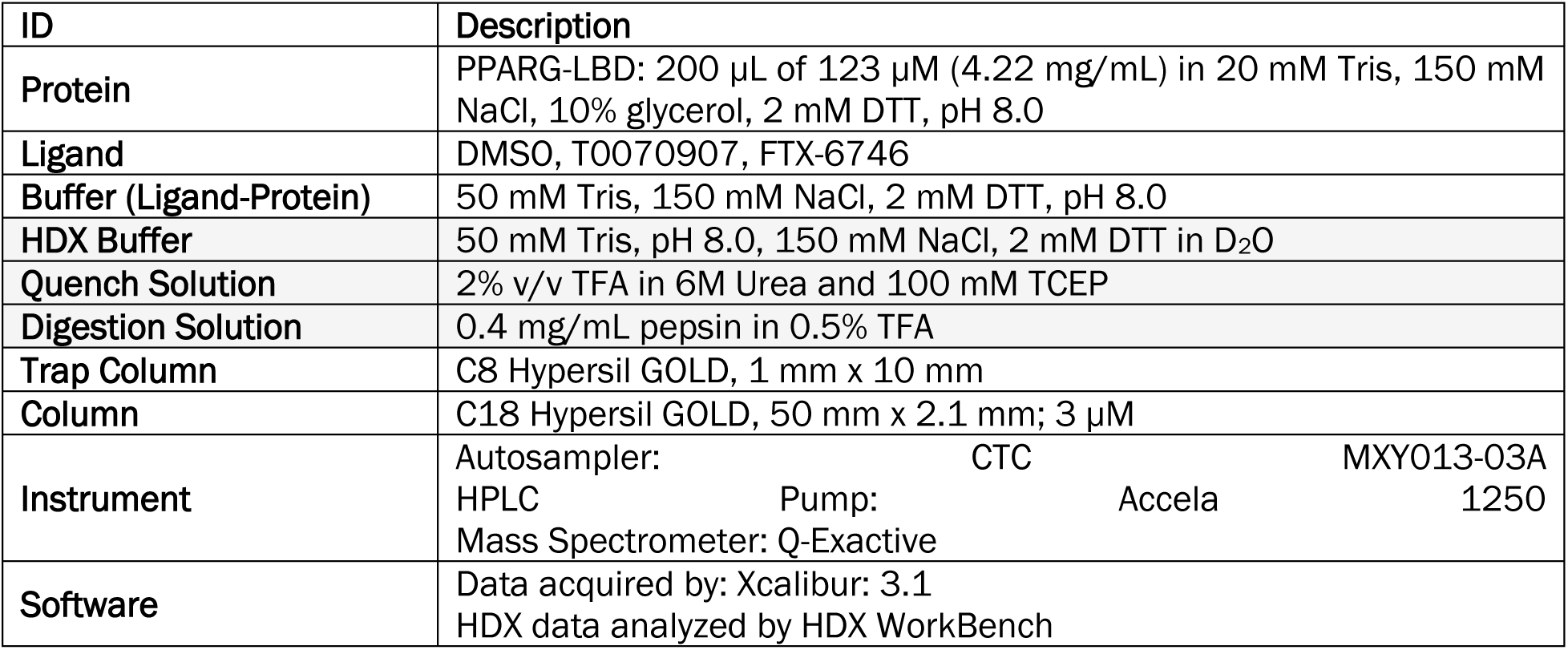
HDX Technical and Buffer Specifications.

## Preparation of FTX-6746, FTX-6713, FTX-6822 and FX-909

### Preparation of FTX-6746, 4-Chloro-3-(5,7-difluoro-4-oxo-1,4-dihydroquinolin-2-yl)benzonitrile

#### Step 1. 2-Chloro-5-cyanobenzoyl chloride

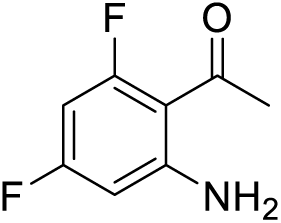

#### Step 1. 1-(2-Amino-4,6-difluorophenyl)ethanone

To a solution of 3,5-difluoroaniline (8.9 g, 68.9 mmol, 1.0 *equiv.*) in CH_3_CN (85 mL) was added BCl_3_ (1 M, 72.4 mL, 1.05 *equiv.*) at 0 °C. Then AlCl_3_ (10.1 g, 75.8 mmol, 4.1 mL, 1.1 *eq*uiv.) was added to the mixture in three portions and the mixture was then stirred at 80 °C for 16 hours. The mixture was cooled to 0 °C and then aqueous HCl (4M, 80 mL) was added and the mixture was stirred at 80 °C for 2 hours. The mixture was cooled to room temperature and extracted with EtOAc (2 x 150 mL). The combined organic layers were washed with saturated aqueous NaHCO_3_ solution (2 x 50 mL), dried over Na_2_SO_4_, filtered, and concentrated under reduced pressure to afford the title compound (8.0 g, 68% yield) as a light-yellow solid.

LCMS: [M+H] for (C_8_H_7_F_2_NO) calculated *m/z* = 172.0, found *m/z* = 172.1.

^1^H NMR (400 MHz, CDCl_3_), δ 6.5 (br s, 2H), 6.0 - 6.2 (m, 2H), 2.6 (d, *J* = 8.4 Hz, 3H).

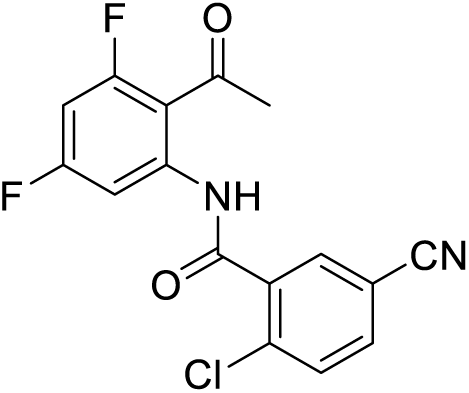

#### Step 2. N-(2-acetyl-3,5-difluorophenyl)-2-chloro-5-cyanobenzamide

A solution of 2-chloro-5-cyano-benzoic acid (2.5 g, 13.8 mmol) in SOCl_2_ (25 mL) was stirred at 80 °C for 1 hour. The reaction mixture was then cooled to room temperature and concentrated under reduced pressure to afford the 2-chloro-5-cyanobenzoyl chloride (2.8 g, crude) as a yellow solid.

To a solution of 1-(2-amino-4,6-difluoro-phenyl)ethanone (2 g, 11.7 mmol, 1.0 *equiv.*) in THF (20 mL) was added NaH (467 mg, 11.7 mmol, 60% dispersion in oil, 1.0 *equiv.*) at 0 °C. The mixture was stirred for 30 minutes before the dropwise addition of a solution of 2-chloro-5-cyano-benzoyl chloride (2.6 g, 12.8 mmol, 1.1 *equiv.*) in THF (10 mL). The mixture was stirred at room temperature for 16 hours. The reaction mixture was quenched by the addition saturated aqueous NH_4_Cl (15 mL) at 15 °C, diluted with water (20 mL), and filtered. The filter cake was triturated with EtOAc (20 mL) and filtered to afford the title compound (2.4 g, 61% yield) as a white solid.

LCMS: [M+H] for (C_16_H_9_F_3_N_2_O_2_) calculated *m/z* = 335.0, found *m/z* = 335.0.

^1^H NMR (400 MHz, DMSO-d_6_): δ 11.2 (s, 1H), 8.1 (d, *J* = 2.0 Hz, 1H), 8.0 (dd, *J* = 8.4, 2.2 Hz, 1H), 7.8 (d, *J* = 8.4 Hz, 1H), 7.5 - 7.5 (m, 1H), 7.3 (ddd, *J* = 11.2, 8.8, 2.2 Hz, 1H), 2.5 - 2.6 (m, 3H).

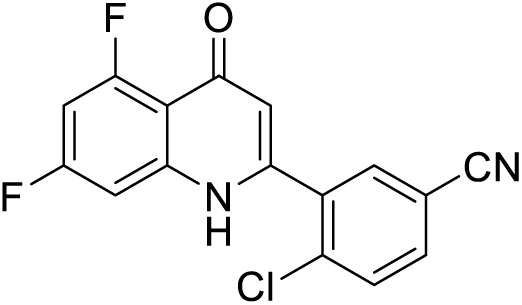

#### Step 3. 4-Chloro-3-(5,7-difluoro-4-oxo-1,4-dihydroquinolin-2-yl)benzonitrile

To a solution of N-(2-acetyl-3,5-difluoro-phenyl)-2-chloro-5-cyano-benzamide (2.5 g, 7.5 mmol, 1.0 *equiv.*) in dioxane (40 mL) was added NaOH (3.0 g, 74.7 mmol, 10.0 *eq*uiv.). The mixture was stirred at 110 °C for 1.5 hours. The pH of the reaction mixture was adjusted to 5 with aqueous HCl (1 M) and then diluted with water (30 mL) and extracted with EtOAc (3 x 100 mL). The combined organic layers were washed with brine (2 x 100 mL), dried over Na_2_SO_4_, filtered, and concentrated under reduced pressure to afford a crude residue that was purified by preparative HPLC (column: Welch Xtimate C18 250 x 70mm x 10um; mobile phase: 15-45% acetonitrile in water (10mM NH_4_HCO_3_)). This afforded the title compound (570 mg, 24% yield, 98% purity) as a white solid after concentration under reduced pressure. LCMS: [M+H] for (C_16_H_7_ClF_2_N_2_O) calculated *m/z* = 317.0, found *m/z* = 317.0.

^1^H NMR (400 MHz, DMSO-d_6_): δ 9.2 - 10.3 (m, 1H), 8.2 (d, *J* = 2.0 Hz, 1H), 8.0 (dd, *J* = 8.4, 2.0 Hz, 1H), 7.9 (d, *J* = 8.4 Hz, 1H), 7.0 - 7.2 (m, 2H), 6.1 (s, 1H).

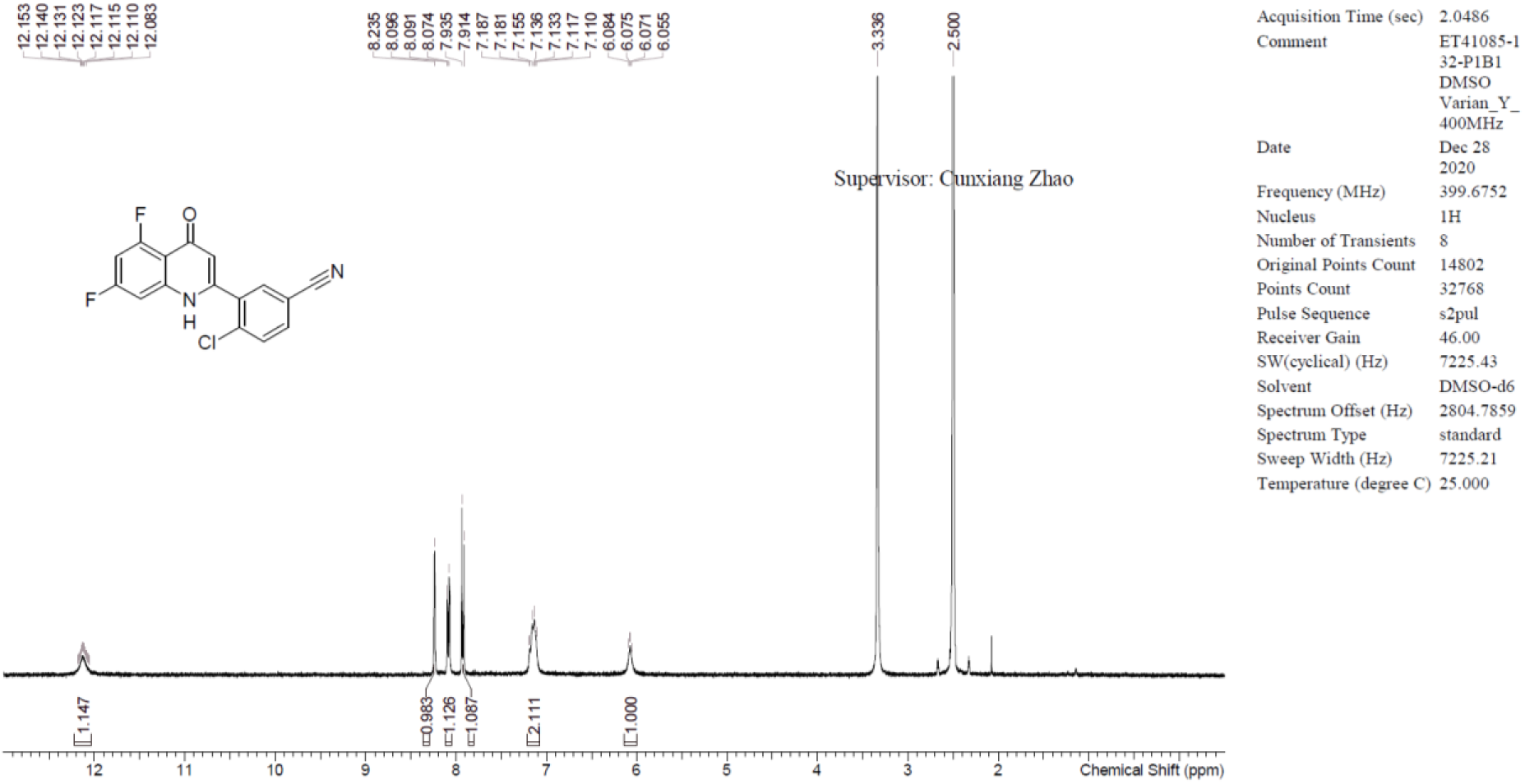

### Preparation of FTX-6822, 3-(5,7-difluoro-4-oxo-1,4-dihydro-2-quinolyl)-4-fluorobenzonitrile

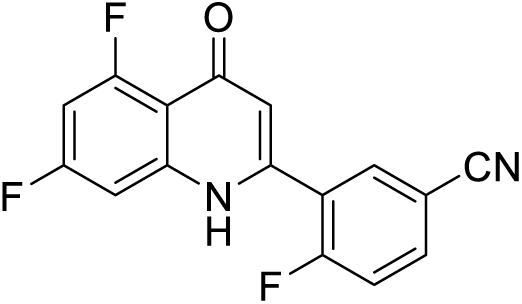

This compound was prepared following the procedure described above for FTX-6746 substituting 5- cyano-2-fluorobenzoic acid for 2-chloro-5-cyano-benzoic acid in step 2.

LCMS: [M+H] for (C_16_H_7_F_3_N_2_O) calculated *m/z* = 300.1, found *m/z* = 300.0.

^1^H NMR (400 MHz, DMSO-d_6_): δ 12.1 (br s, 1H), 8.3 (br d, J = 5.4 Hz, 1H), 8.2 (ddd, J = 8.6, 4.6, 2.0 Hz, 1H), 7.7 (t, J = 9.4 Hz, 1H), 7.1 - 7.3 (m, 2H), 6.1 - 6.4 (m, 1H).

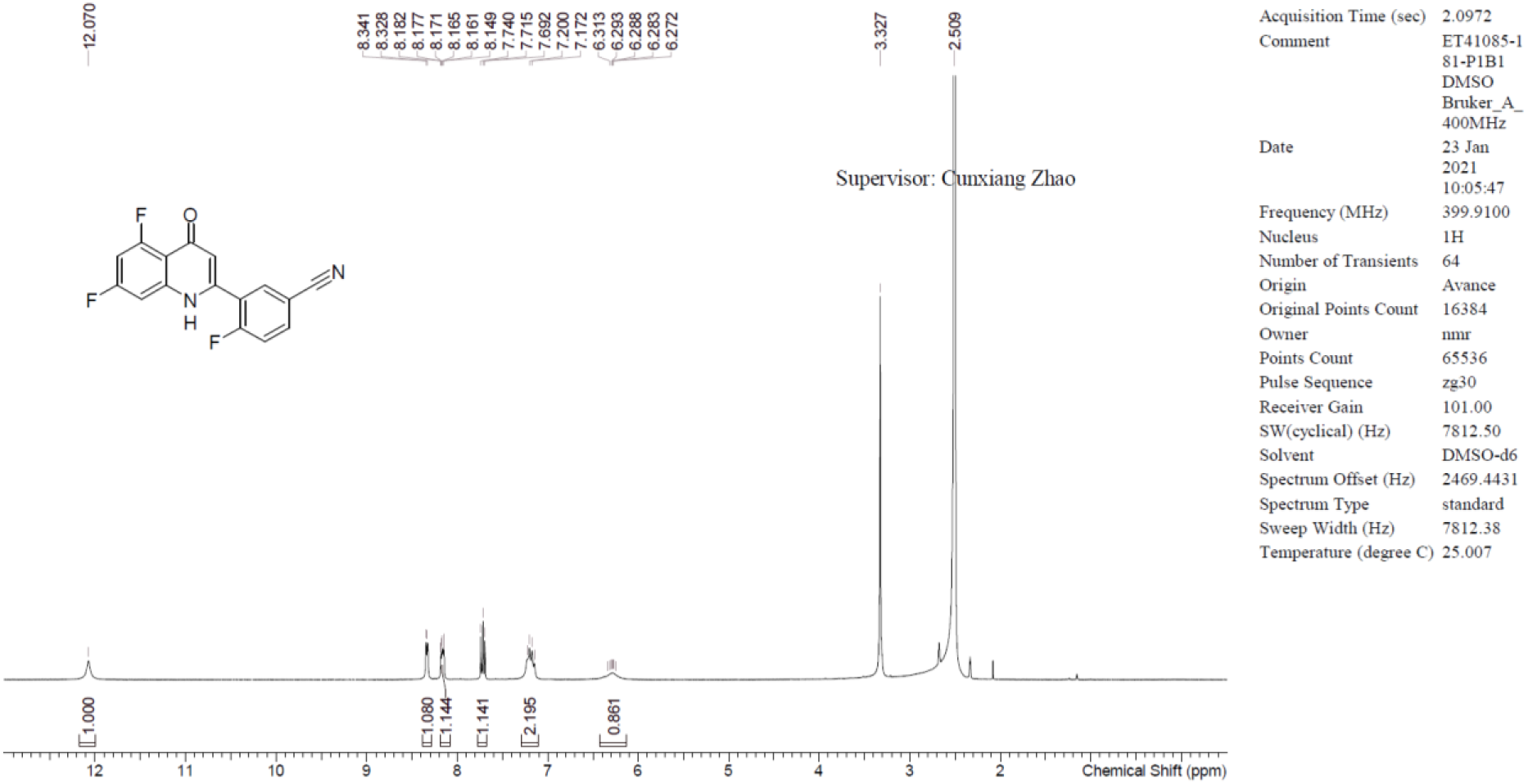

### Preparation of FTX-6713, 4-bromo-3-(5,7-difluoro-4-oxo-1,4-dihydroquinolin-2-yl)benzonitrile

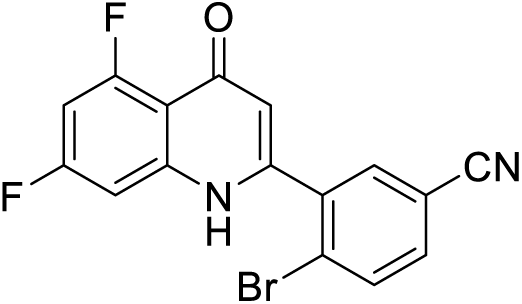

This compound was prepared following the procedure described above for FTX-6746 substituting 5- cyano-2-bromobenzoic acid for 2-chloro-5-cyano-benzoic acid in step 2.

LCMS: [M+H] for (C_16_H_7_BrF_2_N_2_O) calculated *m/z* = 361.0, found *m/z* = 361.0.

^1^H NMR (400 MHz, DMSO-d_6_): δ 8.08 (s, 1H), 8.01 (d, J = 8.4 Hz, 1H), 7.90 (s, 1H), 7.08 (br d, J = 10.2 Hz, 1H), 6.94 - 7.04 (m, 1H), 6.01 (s, 1H).

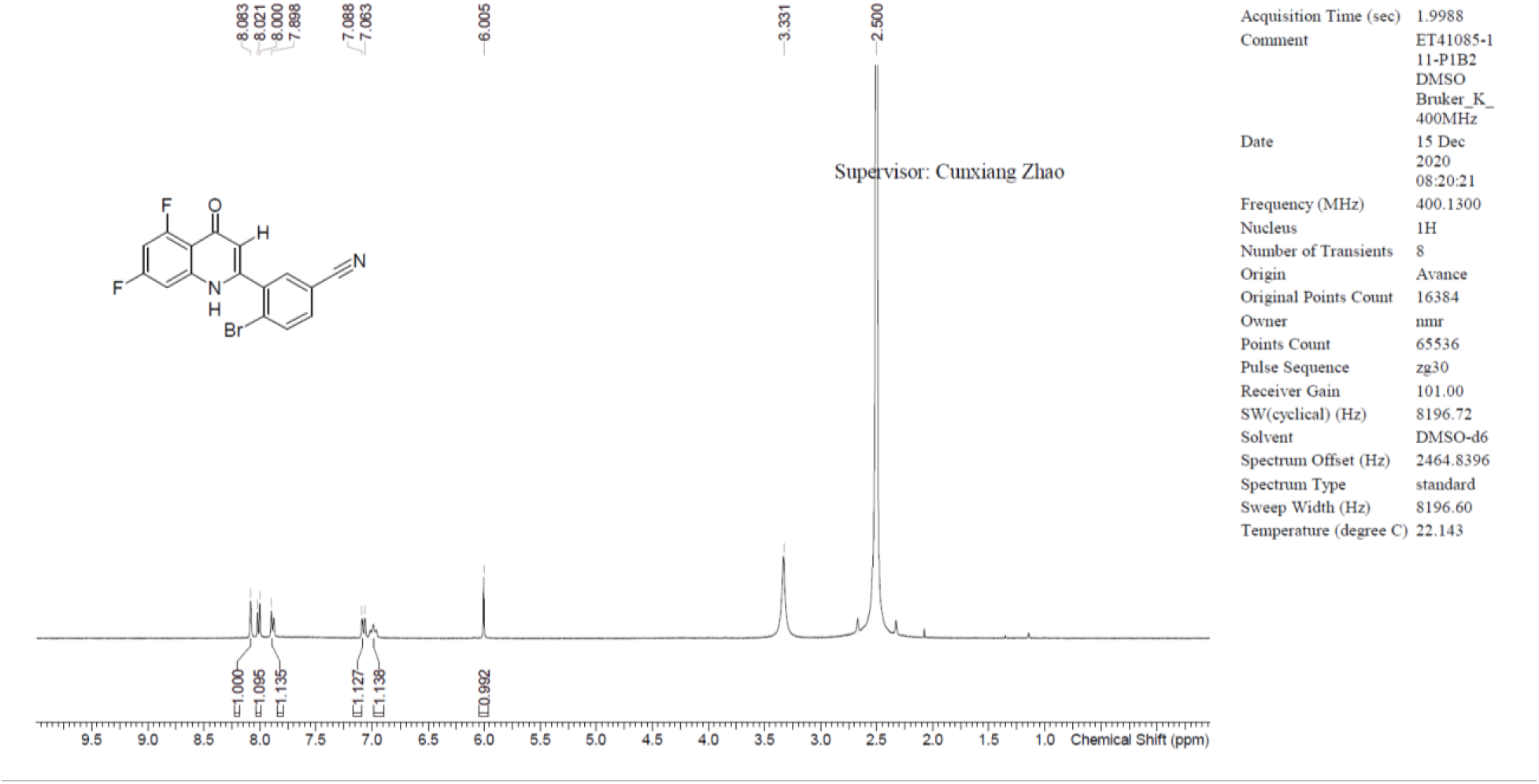

### Preparation of FX-909, 3-(5,7-difluoro-4-oxo-1,4-dihydroquinolin-2-yl)-4-(methylsulfonyl)-benzonitrile

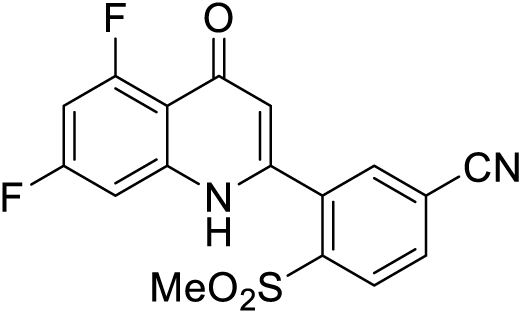

### 3-(5,7-difluoro-4-oxo-1,4-dihydroquinolin-2-yl)-4-(methylsulfonyl)benzonitrile

To a mixture of 4-chloro-3-(5,7-difluoro-4-oxo-1,4-dihydroquinolin-2-yl)benzonitrile, FTX-6746 (100 mg, 316 μmol, 1.0 equiv.) in DMSO (3 mL) were added sodium methanesulfinate (41.9 mg, 411 μmol, equiv.), K_3_PO_4_ (67.0 mg, 316 μmol, 1.0 equiv.), CuI (6.0 mg, 32 μmol, 0.1 equiv.) and quinolin-8-ol (4.6 mg, 32 μmol, 0.1 equiv.) at 20 °C under N_2._ The mixture was stirred at 120°C for 24 hours. The reaction mixture was diluted with H_2_O (30 mL) and extracted with ethyl acetate (3 x 30 mL). The combined organic layers were washed with brine (20 mL), dried with anhydrous Na_2_SO_4_, filtered, and concentrated under vacuum. The residue was purified by preparative HPLC (column: Waters Xbridge BEH C18 100mm x 30mm x 10um; Mobile phase: 25%-55% acetonitrile in water (+NH_4_HCO_3_)) to afford the title compound (37.4 mg, 33% yield, 99.3% purity) as a white solid.

LCMS [M+H] for (C_16_H_7_BrF_2_N_2_O) calculated *m/z* = 361.0, found *m/z* = 361.0. 361.0.

^1^H NMR (400 MHz, METHANOL-d_4_) δ 8.36 (d, *J* = 8.4 Hz, 1H), 8.20 (dd, *J* = 1.6, 8.4 Hz, 1H), 8.13 (d, *J* = 1.4

Hz, 1H), 7.06 (br d, *J* = 9.4 Hz, 1H), 6.99 (ddd, *J* = 2.4, 9.4, 11.8 Hz, 1H), 6.31 (s, 1H), 3.20 (s, 3H).

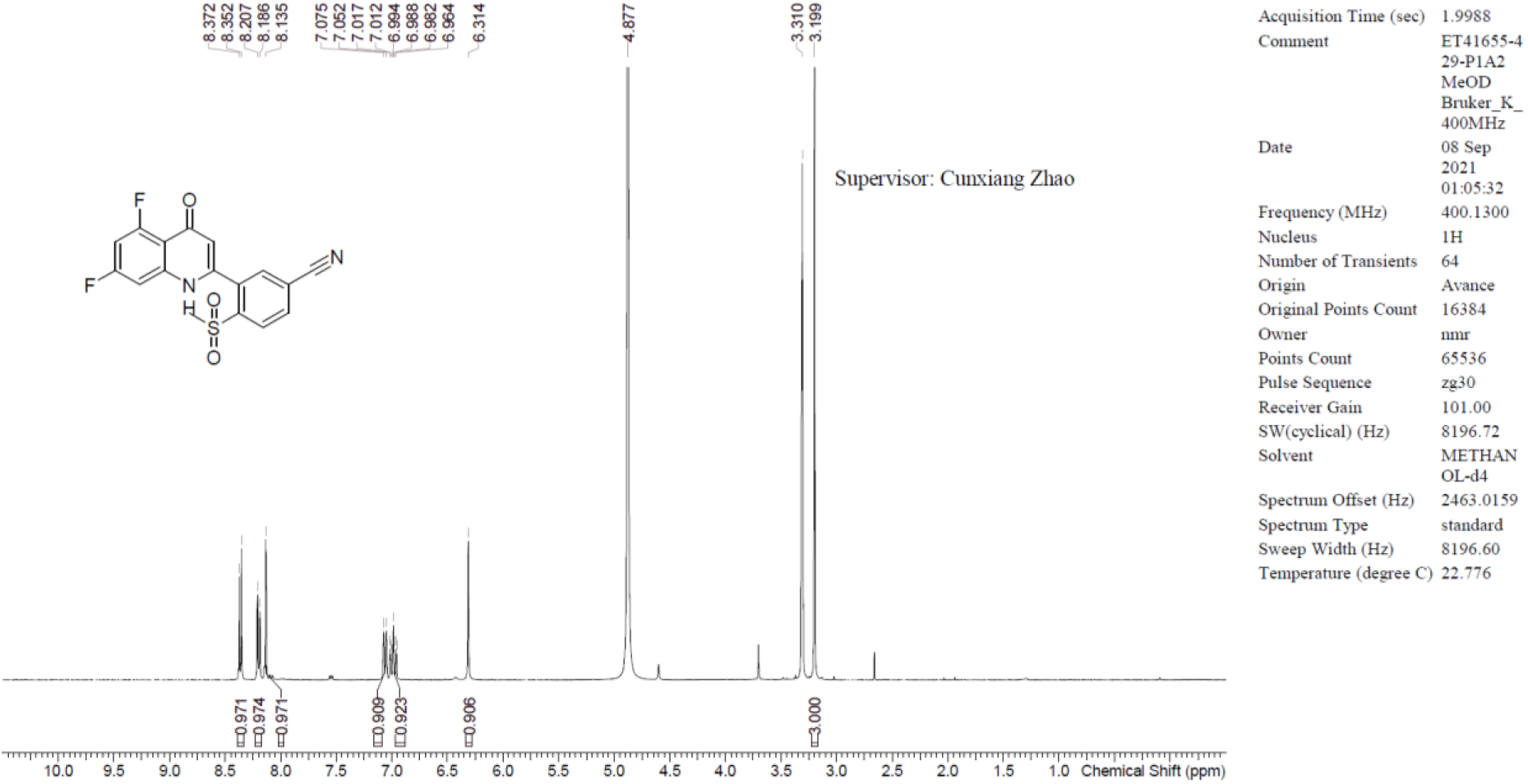

## References

1. Lehrke, M. & Lazar, M. A. The Many Faces of PPARγ. Cell 123, 993–999 (2005).

2. Varley, C. L. et al. Role of PPAR γ and EGFR signalling in the urothelial terminal differentiation programme. J Cell Sci 117, 2029–2036 (2004).

3. Liu, C. et al. Pparg promotes differentiation and regulates mitochondrial gene expression in bladder epithelial cells. Nat Commun 10, 4589 (2019).

4. Chrisman, I. M. et al. Defining a conformational ensemble that directs activation of PPARγ. Nat Commun 9, 1794 (2018).

5. Brust, R. et al. A structural mechanism for directing corepressor-selective inverse agonism of PPARγ. Nat Commun 9, 4687 (2018).

6. Chandra, V. et al. Structure of the intact PPAR-γ–RXR-α nuclear receptor complex on DNA. Nature 456, 350–356 (2008).

7. Pacini, C. et al. Integrated cross-study datasets of genetic dependencies in cancer. Nat. Commun. 12, 1661 (2021).

8. 8. Dempster, J. M., et al. Extracting Biological Insights from the Project Achilles Genome-Scale CRISPR Screens in Cancer Cell Lines. bioRxiv 720243 (2019) doi:10.1101/720243.

9. 9. Dempster, J. M. et al. Chronos: a CRISPR cell population dynamics model. Biorxiv 2021.02.25.432728 (2021) doi:10.1101/2021.02.25.432728.

10. Meyers, R. M. et al. Computational correction of copy number effect improves specificity of CRISPR–Cas9 essentiality screens in cancer cells. Nat. Genet. 49, 1779–1784 (2017).

11. Goldstein, J. T. et al. Genomic activation of PPARG reveals a candidate therapeutic axis in bladder cancer. Cancer Res 77, canres.1701.2017 (2017).

12. Rochel, N. et al. Recurrent activating mutations of PPARγ associated with luminal bladder tumors. Nat Commun 10, 253 (2019).

13. Korpal, M. et al. Evasion of immunosurveillance by genomic alterations of PPARγ/RXRα in bladder cancer. Nat Commun 8, 103 (2017).

14. Irwin, S. et al. Biochemical and structural basis for the pharmacological inhibition of nuclear hormone receptor PPARγ by inverse agonists. J. Biol. Chem. 298, 102539 (2022).

15. Lee, G. et al. T0070907, a Selective Ligand for Peroxisome Proliferator-activated Receptor γ, Functions as an Antagonist of Biochemical and Cellular Activities*. J Biol Chem 277, 19649– 19657 (2002).

16. Shang, J. et al. A molecular switch regulating transcriptional repression and activation of PPARγ. Nat Commun 11, 956 (2020).

17. Orsi, D. L. et al. Discovery and characterization of orally bioavailable 4-chloro-6- fluoroisophthalamides as covalent PPARG inverse-agonists. Bioorgan Med Chem 117130 (2022) doi:10.1016/j.bmc.2022.117130.

18. Orsi, D. L. et al. Discovery and Structure-Based Design of Potent Covalent PPARγ Inverse- Agonists BAY-4931 and BAY-0069. J Med Chem (2022) doi:10.1021/acs.jmedchem.2c01379.

19. Hu, X. & Lazar, M. A. The CoRNR motif controls the recruitment of corepressors by nuclear hormone receptors. Nature 402, 93–96 (1999).

20. Kilu, W., Merk, D., Steinhilber, D., Proschak, E. & Heering, J. Heterodimer formation with retinoic acid receptor RXRα modulates coactivator recruitment by peroxisome proliferator-activated receptor PPARγ. J. Biol. Chem. 297, 100814 (2021).

21. Freitag, C. M. & Miller, R. J. Peroxisome proliferator-activated receptor agonists modulate neuropathic pain: a link to chemokines? Front. Cell. Neurosci. 8, 238 (2014).

22. Miyamae, Y. Insights into Dynamic Mechanism of Ligand Binding to Peroxisome Proliferator-Activated Receptor γ toward Potential Pharmacological Applications. Biol. Pharm. Bull. 44, 1185–1195 (2021).

23. Mosure, S. A. et al. Structural Basis of Altered Potency and Efficacy Displayed by a Major in Vivo Metabolite of the Antidiabetic PPARγ Drug Pioglitazone. J. Med. Chem. 62, 2008–2023 (2019).

24. Irwin, S. et al. Biochemical and structural basis for the pharmacological inhibition of nuclear hormone receptor PPARγ by inverse agonists. J. Biol. Chem. 298, 102539 (2022).

25. Leesnitzer, L. M. et al. Functional Consequences of Cysteine Modification in the Ligand Binding Sites of Peroxisome Proliferator Activated Receptors by GW9662. Biochemistry-us 41, 6640–6650 (2002).

26. Kwan, E. E., Zeng, Y., Besser, H. A. & Jacobsen, E. N. Concerted nucleophilic aromatic substitutions. Nat. Chem. 10, 917–923 (2018).

27. Duan, S. Z. et al. Hypotension, lipodystrophy, and insulin resistance in generalized PPARγ- deficient mice rescued from embryonic lethality. J Clin Invest 117, 812–822 (2007).

28. Gilardi, F. et al. Systemic PPARγ deletion in mice provokes lipoatrophy, organomegaly, severe type 2 diabetes and metabolic inflexibility. Metabolis 95, 8–20 (2019).

29. Marciano, D. P. et al. Pharmacological repression of PPARγ promotes osteogenesis. Nat. Commun. 6, 7443 (2015).

30. Beausoleil, S. A., Villén, J., Gerber, S. A., Rush, J. & Gygi, S. P. A probability-based approach for high-throughput protein phosphorylation analysis and site localization. Nat. Biotechnol. 24, 1285–1292 (2006).

31. Huttlin, E. L. et al. A Tissue-Specific Atlas of Mouse Protein Phosphorylation and Expression. Cell 143, 1174–1189 (2010).

32. McAlister, G. C. et al. Increasing the Multiplexing Capacity of TMTs Using Reporter Ion Isotopologues with Isobaric Masses. Anal. Chem. 84, 7469–7478 (2012).

33. Elias, J. E. & Gygi, S. P. Target-decoy search strategy for increased confidence in large-scale protein identifications by mass spectrometry. Nat. Methods 4, 207–214 (2007).

34. Vestal, B. E., Wynn, E. & Moore, C. M. lmerSeq: an R package for analyzing transformed RNA-Seq data with linear mixed effects models. BMC Bioinform. 23, 489 (2022).

35. Korotkevich, G. et al. Fast gene set enrichment analysis. bioRxiv 060012 (2021) doi:10.1101/060012.

36. Biton, A. et al. Independent Component Analysis Uncovers the Landscape of the Bladder Tumor Transcriptome and Reveals Insights into Luminal and Basal Subtypes. Cell Reports 9, 1235–1245 (2014).

37. Love, M. I., Huber, W. & Anders, S. Moderated estimation of fold change and dispersion for RNA-seq data with DESeq2. Genome Biol. 15, 550 (2014).

38. Vivian, J. et al. Toil enables reproducible, open source, big biomedical data analyses. Nat. Biotechnol. 35, 314–316 (2017).

39. 39. DepMap, B. DepMap 24Q2 Public. Figshare+. Dataset. 10.25452/figshare.plus.25880521.v1 (2024).

